# Mitochondria-containing large extracellular vesicles target mouse motor neurons upon intramuscular injection

**DOI:** 10.1101/2025.09.15.675842

**Authors:** Paromita Paul Pinky, Zhong-Min Wang, Purva Khare, Jhanvi R. Jhaveri, Vivek Basudkar, Krithika S. Rao, Audrey Lawrence, Adithri Pingali, Kandarp M. Dave, Donna B. Stolz, Ming Sun, Si-yang Zheng, Sruti S. Shiva, Carol Milligan, Osvaldo Delbono, Devika S Manickam

## Abstract

Amyotrophic Lateral Sclerosis (**ALS**) is a neurological disorder that causes progressive degeneration of motor neurons. Mitochondrial dysfunction accelerates neurodegeneration aggravating the severity of ALS. We hypothesized that increasing the mitochondrial function of motor neurons may promote neuronal survival. Therefore, we investigated the potential of neuron-derived mitochondria containing extracellular vehicles (**EVs**) as a novel therapeutic approach for ALS using differentiated NSC-34 cells as a surrogate for neurons. Neuron derived-large EVs (**lEVs**) but not small EVs (**sEVs**) contained mitochondria. However, we observed increased cell viability and oxygen consumption rates in heat-stressed neurons treated with both sEVs and lEVs suggesting improved mitochondrial function in recipient neurons. The increased oxygen consumption rates in sEV-treated heat-stressed neurons was accompanied by a *greater proton leak* compared to lEV treatment. The greater proton leak observed with sEVs likely suggests a lower efficiency of oxidative phosphorylation compared to that achieved by cells treated with mitochondria-containing lEVs. These findings suggest that mitochondrial components present in sEVs, such as proteins and mitochondrial DNA, may too contribute to improving cellular respiration. Furthermore, we have demonstrated that lEV mitochondria are transported into the lumbar spinal cord motor neurons following intramuscular injection in C57BL/6 mice in a EV dose-dependent manner. Collectively, for the first time, we have demonstrated the therapeutic effects of neuronal EVs in recipient heat-stressed neurons and the delivery of lEV mitochondria to spinal cord motor neurons *in vivo* without any EV surface modifications for neuronal targeting. Further studies will determine the therapeutic efficacy of mitochondria-containing EVs in the SOD1^G93A^ transgenic mouse model of ALS.

**Graphical abstract:** 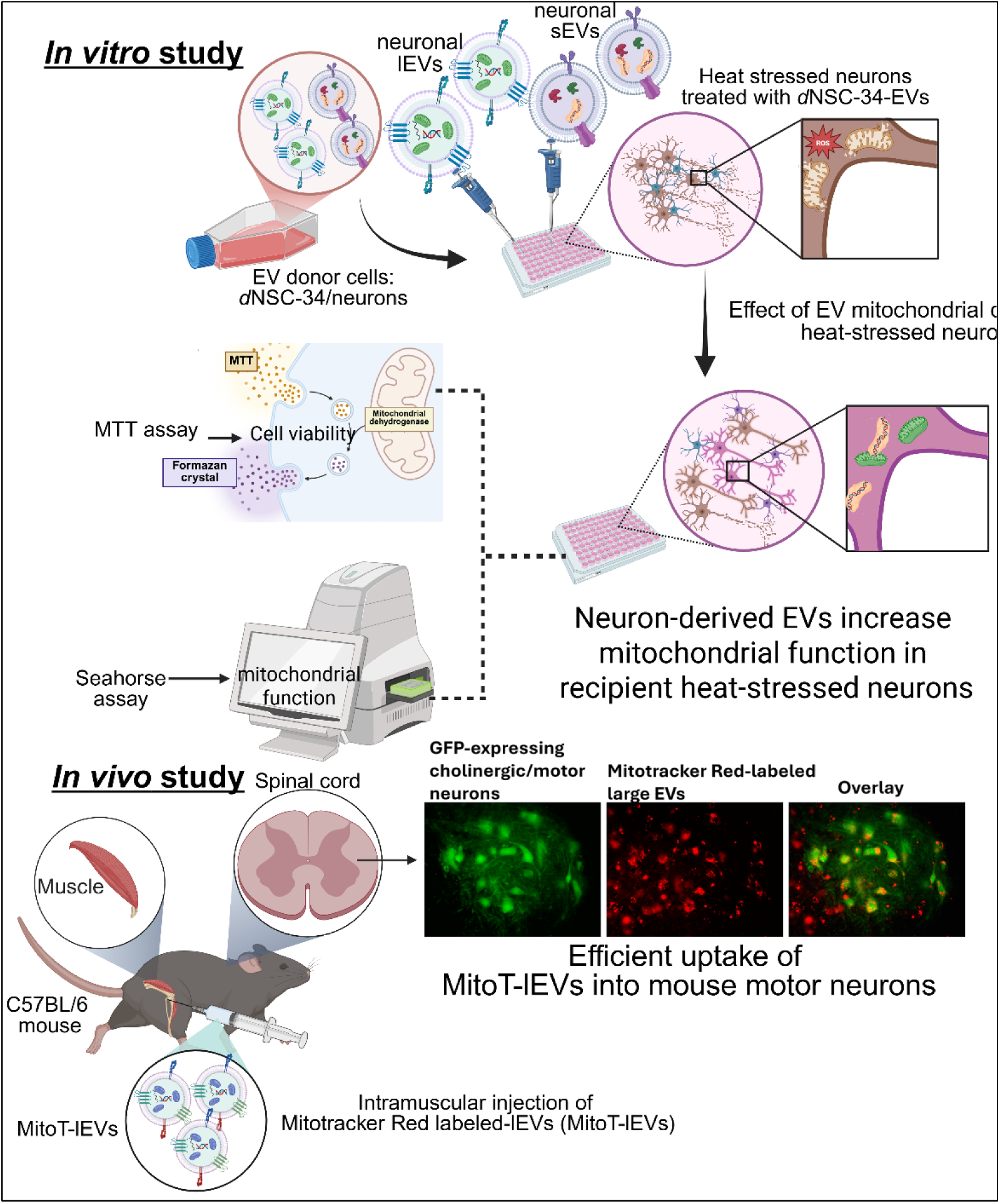

## 1. Introduction

ALS is a neurodegenerative disorder characterized by the degeneration of upper motor neurons in the motor cortex and lower motor neurons in the brain stem and the spinal cord [2]. ALS can be sporadic or familial in origin, however, both these types are clinically indistinguishable [3]. Patients experience muscle weakness, progressive motor disability, and eventually respiratory arrest or an associated infection leading to death. In most ALS cases, the average expected lifespan is only a dismal five years post-diagnosis [3]. In 1869, Charcot first described ALS using clinical cases and autopsy [4]. Despite the early identification of ALS, the pathophysiology of ALS is unclear. Commonly reported ALS pathology involves glutamate excitotoxicity, oxidative stress, mitochondrial dysfunction, axonal transport abnormalities, apoptosis, neuroinflammation, RNA and protein metabolism [2, 3, 5, 6]. Among these pathophysiological events, mitochondrial dysfunction and its structural abnormalities mark the early features of motor neuron degeneration and have been extensively documented in *vitro, in vivo,* and in postmortem samples from ALS patients [7–11]. Collectively, these reports have revealed that structural and functional abnormalities of mitochondria result in increased oxidative stress (due to increased generation of reactive oxygen species), reduced cellular respiration and ATP production, disrupted calcium homeostasis, and decreased mitochondrial membrane potential. Consequently, motor neurons undergo progressive dysfunction and eventual physical degeneration which aggravates the severity of ALS, shortening the life expectancy of affected patients.

Currently, the FDA-approved drugs for ALS treatment can extend the life span of ALS patients only by several months. Riluzole (the first FDA-approved drug to treat ALS) inhibits glutamate-induced excitotoxicity and decreases intracellular calcium concentration likely leading to the neuroprotective effect [12]. Tofersen is an antisense oligonucleotide that targets the mutation in the superoxide dismutase 1 (**SOD1**) gene [13]. Edaravone is a scavenger of reactive oxygen species (**ROS**) leading to cell survival [14] [15]. Dextromethorphan HBr and quinidine sulfate have been used as an off-label treatment in ALS patients [16, 17]. This combination works to treat the pseudobulbar effect, a secondary effect in many neurological conditions [18]. Relyvrio is a combination of sodium phenylbutyrate and taurursodiol and was recently approved by the FDA in 2022 to treat ALS. The mechanism of action involved reduction of endoplasmic reticulum stress and mitochondrial dysfunction, the two major contributors to neurodegeneration. However, it was voluntarily withdrawn from the market as of April 2024 based on topline results from the phase 3 trial due to lack of efficiency [18]. Several drugs (dexpramipexole, minocycline, olesoxime, *etc*.) targeting mitochondrial function and ROS were found inefficacious during clinical trials [19, 20]. As these mitochondria therapies only target individual events of mitochondrial dysfunction such as scavenging ROS or restoring calcium homeostasis individually, they may not be sufficient to improve mitochondrial function and ultimately, motor neuron survival. Thus, there is an unmet clinical need to develop new therapeutics to slow down the progression of ALS thereby prolonging the life span of the patients.

Based on our prior works where we have demonstrated the transfer of mitochondria via extracellular vesicles (**EVs**) into recipient brain endothelial cells (**BECs**) and brain slice neurons [21–24], we hypothesized that EVs derived from neurons may exhibit similar effects in an experimental model of ALS. Previously, we isolated, characterized, and determined the protective effects of BEC-derived EVs in recipient oxygen-glucose-deprived BECs and in a mouse model of ischemic stroke [21–24]. Therefore, in this study, we investigated whether the delivery of functional mitochondria via neuron-derived EVs may increase mitochondrial function in heat-stressed neurons, an experimental model of ALS. In addition to our prior works, other published reports have also demonstrated that intercellular mitochondrial transfer rescues and revitalizes tissue in various pathologies, including central nervous system disorders [25, 26], respiratory system disorders [27, 28], and musculoskeletal system disorders [29]. Tunneling nanotubes and EV-mediated mitochondrial transfer are the reported mechanisms of cell-to-cell mitochondrial transfer [30, 31].

EVs are cell-derived heterogeneous particles secreted into extracellular spaces under physiological and pathological conditions. EVs carry an array of bioactive cargo enclosed within their lipid membranes. Among several subtypes of EVs, two major EV subtypes distinguished based on their particle diameters [32] referred to as large **EVs (**lEVs >200 nm particle diameter) and small EVs **(sEVs** <200 nm particle diameter) are being studied extensively because of their promising therapeutic potential in various pathological conditions. Mounting evidence reveals the presence of mitochondria and mitochondrial DNA in EVs [31, 33–40]. Our lab has previously shown that BEC-derived lEVs but not sEVs contain mitochondria [41, 42]. In contrast, sEVs do not contain mitochondria but carry mitochondrial DNA [34] and mitochondrial proteins [34, 41, 42]. We demonstrated that the delivery of mitochondria containing lEVs improved mitochondrial function and the intracellular ATP level of recipient BECs under oxygen glucose-deprived conditions, an *in vitro* stroke model [42]. Intravenously injected lEVs caused a 50% reduction in infarct volume stroke and also showed a 33% improvement in neurological functions in a mouse model of transient ischemic stroke [24].

Other published works have reported the functional effects of lEV-mediated mitochondria in different pathological conditions. For instance, Islam *et al.* reported that lEV-mediated mitochondria transfer from bone-marrow-derived stromal cells increased cellular ATP levels *in vitro* in recipient alveolar epithelial cells and protected against acute lung injury *in vivo* [43]. O’Brien and workers demonstrated that lEV-treated doxorubicin-injured cardiomyocytes showed improved ATP production, mitochondrial biogenesis, and attenuated apoptosis. These outcomes were attributed to the presence of mitochondria in lEVs [44]. Collectively, these findings revealed that lEVs can successfully transfer their innate mitochondrial content to target recipient cells and exert functional effects by increasing cellular respiration and ATP levels. Furthermore, our previous study showed that recipient BECs preferentially internalize BEC-derived-lEVs compared to macrophage-derived lEVs [45].

EVs are considered as a safer therapeutic approach compared to whole cell therapy. Current evidence demonstrates that only a small fraction of injected stem cells can reach the affected site for proliferation and differentiation which may reduce the expected therapeutic efficacy of stem cell therapy in clinical settings [46]. Furthermore, because of their larger size, stem cells are not able to cross the blood-brain barrier limiting their capability to provide maximum therapeutic benefits [47]. Moreover, the high proliferative capacity of stem cells can have significant oncogenic potential with higher chances of immune rejection by the recipient immune system [3, 48]. Besides, vascular obstruction is a serious concern for cell therapy which can restrict its therapeutic potential [48]. In contrast, EVs are a non-cell based therapeutic approach which can minimize complications associated with whole cell therapy. Furthermore, as cell-derived carriers, EVs tend to show several advantages including reduced immune response, biocompatibility and stability in biological fluids [49]. Therefore, we chose to derive mitochondria-containing lEVs from a neuronal source to investigate their effects in an experimental model of ALS. Considering this scientific premise, we *hypothesize* that the delivery of functional mitochondria containing neuron-derived lEVs will increase mitochondrial function in the heat-stressed neurons resulting in increased cell survival.

We selected the NSC-34 cell line as a surrogate of neurons. NSC-34 is a hybrid cell line comprised of motor neuron-enriched embryonic mouse spinal cord cells fused with neuroblastoma. These cells express neuron-like properties including choline acetyltransferase, acetylcholine synthesis, storage, and release [50]. These cells were experimentally stressed to mimic the pathophysiological events of ALS. In this study, we isolated neuron-derived-lEVs and sEVs using sequential ultracentrifugation [42]. EVs were characterized by measuring their particle size, zeta potential, and dispersity index. lEVs were pelleted down at 20,000×g showed average particle diameters of >200 nm whereas sEV pellets obtained at 120,000×g showed average diameters of <200nm. Cross-sectioned TEM images revealed that lEVs contained mitochondria in their lumen. We performed flow cytometry and fluorescence microscopy to determine lEV-mediated mitochondrial transfer into recipient neurons. MTT and Seahorse assays revealed that both lEVs and sEVs increased cell viability as well as mitochondrial respiration in recipient neurons. A pilot *in vivo* study in C57BL/6 mice demonstrated EV dose-dependent delivery of mitochondria-containing large EVs into spinal cord motor neurons upon intramuscular injection—demonstrating the safety and targetability of delivery to motor neurons. Overall, our results reveal the therapeutic potential of neuron-derived EVs as a new approach to treat ALS.

## 2. Materials and methods

### 2.1. Materials

Mouse motor neuron-like neuroblastoma NSC-34 cells were obtained from CELLutions biosystems (Burlington, Canada). A mouse brain endothelial cell line (bEnd.3, cat# CRL2299) was purchased from ATCC (Manassas, VA). Pierce BCA, Micro BCA protein assay kit, and L-glutamine were purchased from Thermo Scientific (Rockford, IL). Collagen Type I was purchased from Corning (Discovery Labware Inc., Bedford, MA). Neurobasal medium, DMEM/Glutamax medium, and Penicillin-Streptomycin (Pen-Strep) solution were purchased from Gibco (Grand Island, NY). Heat-inactivated fetal bovine serum (FBS) was purchased from HyClone Laboratories (Logan, UT). MitoTracker Deep Red FM was purchased from Invitrogen (Carlsbad, CA). RIPA (5*x*) buffer was purchased from Alfa Aesar (Ward Hill, MA). Rabbit anti-ARF6, rabbit anti-CD9 and Recombinant Alexa Fluor 555 Anti-beta III Tubulin primary antibodies were purchased from Abcam (Boston, MA). Goat anti-rabbit secondary antibody AF790 was purchased from Jackson Immuno Research Lab Inc. (West Grove, PA). 10% Mini-PROTEAN TGX Precast Protein Gels were purchased from Bio-Rad Laboratories, Inc. (Hercules, CA). Oligomycin A was purchased from Sigma-Aldrich (Saint Louis, MO). MTT reagent was purchased from Invitrogen. Polycarbonate centrifuge tubes were purchased from Beckman Coulter, Inc. (Brea, CA). Two-well chamber slides were purchased from Thermo Fisher Scientific (Thermo Scientific Nunc Lab-Tek II Chamber Sl System). Low-volume disposable cuvettes (Malvern ZEN0040) were purchased from Fisher Scientific (Waltham, MA).

### 2.2. Cell culture

#### 2.2.1. NSC-34

NSC-34 cells were maintained in tissue culture flasks or 96-well plates in a humidified 5% CO_2_ incubator at 37 ± 0.5°C (Isotemp, Thermo Fisher Scientific). The cells were cultured in a complete growth medium composed of DMEM high glucose medium supplemented with 10% FBS and 1% Pen-Strep. The cells were supplemented with fresh growth medium every other day until the cells reached 90-95% confluency. NSC-34 cells were maintained between passage (P) numbers P29 and P40 and were used in all experiments. Basal NSC-34 was differentiated into neurons for the isolation of lEVs and sEVs. To differentiate the NSC-34 into neurons, the growth medium was switched to neurobasal (with 4 mM glutamine and 1% Pen-Strep) medium when the NSC-34 cells were at ∼80-85% cell confluency [51]. After switching to the Neurobasal differentiation medium, cell morphology was regularly observed for signs of neurite outgrowth under an EVOS microscope (EVOS FL, Invitrogen).

#### 2.2.2. Mouse brain endothelial (b.End3) cells

Mouse brain endothelial (**b.End3**) cells were maintained in tissue culture flasks in a humidified 5% CO_2_ incubator at 37 ± 0.5°C (Isotemp, Thermo Fisher Scientific). The cells were cultured in complete growth medium composed of DMEM high glucose medium supplemented with 10% FBS and 1% Penicillin-Streptomycin. The cells were supplemented with fresh growth medium every other day until the cells reached 90-95% confluency. b.End3 cells maintained between passage (P) numbers P29 and P40 and were used in all experiments.

### 2.3. Immunostaining of neurite processes of *d*NSC-34 cells

Neurite outgrowths of *d*NSC-34 cells (or control NSC-34 cells) were stained using class III beta-tubulin neuronal marker antibody using the protocol adapted from [51]. NSC-34 cells were seeded at a density of 60,000 cells/chamber in a two-chamber slide. The chamber slide was coated with collagen type-I for 1 hour followed by a PBS wash. Cells were suspended in a Neurobasal differentiation medium after dissociating with TrypLE and added to the chamber slide at the required cell density (for NSC-34 cells, the same protocol was used except that the cells were stained when they reached 80-85% confluency instead of switching them to Neurobasal medium). The cells were cultured for four days and supplemented with fresh differentiation medium on day 2. Post incubation, the cells were fixed with 2% paraformaldehyde in PBS for 30 minutes at room temperature and washed twice with PBS. Cells were permeabilized with 0.1% Triton X-100 in 1*x* PBS for five minutes at room temperature. Post-permeabilization, cells were washed with PBS and incubated with 1:1 LiCOR blocking solution: PBS-T (0.1% Tween 20 in PBS) for one hour at room temperature to block the non-specific binding sites. Cells were then incubated with recombinant Alexa Fluor 555 Anti-beta III Tubulin antibody (1:500 dilution) overnight at 4°C, washed thrice in PBS, and incubated with 10µg/mL Hoechst to label the nucleus. NSC-34 cells and *d*NSC-34 cells were then imaged using an EVOS and Olympus IX 73 epifluorescence inverted microscope (Olympus, Pittsburgh, PA) on an RFP channel to detect beta III tubulin signals and a DAPI channel to detect Hoechst signals at 20X magnification. The morphology of NSC-34 and *d*NSC-34 cells were compared in terms of neurite out-growth.

### 2.4. Isolation of NSC-34 EVs and *d*NSC-34 EVs

A differential ultracentrifugation technique was used to isolate NSC-34-derived EVs and *d*NSC-34-derived EVs (sEVs and lEVs) from the EV-conditioned medium using methods we have previously described [22, 24, 41, 42, 52, 53]. Briefly, NSC-34 cells were cultured in 182 cm^2^ growth area (T175) tissue culture flasks until they reached 90-95% confluency. After that, the cells were washed with PBS and incubated with serum-free medium for 48 h in a humidified CO_2_ incubator at 37 ± 0.5°C.

To isolate *d*NSC-34 EVs, NSC-34 cells were switched to the neurobasal differentiation medium and incubated for 72 hours under a humidified incubator to allow differentiation. After 72 hours of incubation, cells were supplemented with fresh Neurobasal differentiation medium and incubated for 48 hours. The same isolation protocol was followed to isolate NSC-34 EVs and *d*NSC-34 EVs. Post-incubation, the conditioned medium was collected in a polypropylene centrifuge tube and centrifuged at 300×g for 11 minutes at 4^°^C to remove all dead cells using a Sorvall ST 8R centrifuge (ThermoFisher Scientific, Osterode am Harz, Germany). Subsequently, the collected supernatant was further centrifuged at 2000 ×g for 22 min at 4°C to pellet down apoptotic bodies and cell debris using a Sorvall ST 8R centrifuge (ThermoFisher Scientific, Osterode am Harz, Germany). Following this, the supernatant was transferred into polycarbonate tubes and centrifuged at 20,000 ×g for 45 min at 4°C to pellet down lEVs using an Optima XE-90 ultracentrifuge fitted with a 50.2 Ti rotor (Beckman Coulter, Indianapolis, IN). Next, the supernatant was collected and filtered through a 0.22 µm Millipore express (PES) membrane syringe filter and centrifuged at 120,000 ×g for 70 min at 4°C to collect sEVs. lEV pellets and sEV pellets were suspended in PBS and stored at -80°C until further use.

### 2.5. MicroBCA Assay

Total EV protein content was measured using Pierce MicroBCA assay, where the EV samples were lysed using 1*x* RIPA buffer at 1:32 dilution (v/v). Next, 150 µL of the lysed EV samples and BSA standards (0.5–200 μg/mL) were pipetted in the 96-well plate and mixed with an equal volume of the Micro BCA working reagent (reagent A: reagent B: reagent C at 25:24:1 volume ratio). The plate was then incubated at 37°C for 2 h. Post incubation, the absorbance was measured at 562 nm using a SYNERGY HTX multi-mode reader (BioTek Instruments Inc., Winooski, VT).

### 2.6. Dynamic light scattering (DLS)

Particle diameter, dispersity index and zeta potential of lEVs and sEVs were measured using DLS on a Malvern Zetasizer Pro (Malvern Panalytical Inc., Westborough, PA). Sample were prepared at 0.1 mg protein/mL concentration by suspending in PBS and 10 mM HEPES buffer at pH 7.4 for particle size and zeta potential measurements, respectively. Average particle diameter, dispersity index, and zeta potential values were reported as mean ± standard deviation of n=3

### 2.7. Nanoparticle tracking analysis (NTA)

EV particle concentrations (particle/ml) were determined on a multiple-laser ZetaView f-NTA Nanoparticle Tracking Analyzer (Particle Metrix Inc., Mebane, NC). For NTA, stock samples of lEVs and sEVs were diluted in PBS before analysis. Five 60 s videos were acquired at 520 nm laser wavelength for concentration measurements. Data were reported as mean ± standard deviation.

### 2.8. Transmission Electron Microscopy (TEM) of EVs

#### 2.8.1. Negative-stain images of EVs

The EV samples diluted in PBS were mounted on formvar-coated copper grids for 5 minutes and negatively stained with 1% uranyl acetate. After staining, specimens were examined on a JEOL-1400 Flash transmission electron microscope (JEOL, Peabody, 268 MA, USA) fitted with a bottom mount AMT Biosprint 12, 12mp digital camera (Advanced Microscopy Techniques, Danvers, MA) [42].

#### 2.8.2. TEM image of cross-sectioned EVs

For capturing the cross-sectioned TEM images of lEVs and sEVs, samples were processed following the protocol that was previously described by us [24, 42]. Suspension of EVs was pelleted at 100,000×g in a Beckman Airfuge for 20 min followed by fixation with 2.5% glutaraldehyde in PBS overnight. After removing the supernatant, pellets were washed thrice with PBS and post-fixed in 1% OsO_4_, and 1% K_3_Fe (CN)_6_ for 1 h. After three additional PBS washes, a graded series of 30–100% ethanol was used to dehydrate the pellet. After several changes of 100% resin over 24 h, the pellet was embedded in a final change of resin, cured at 37 ◦C overnight, followed by additional hardening at 65 ◦C for two more days. Ultrathin (70 nm) sections were collected on 200 mesh copper grids, stained with 2% uranyl acetate in 50% methanol for 10 min, followed by 1% lead citrate for seven min. Sections were imaged using a JEOL JEM 1400 Plus transmission electron microscope (Peabody, MA) at 80 kV fitted with a side mount AMT 2 k digital camera (Advanced Microscopy Techniques, Danvers, MA) [42].

### 2.9. Western blotting

Characteristic protein markers of EVs isolated from NSC-34 and *d*NSC-34 cells were detected using western blotting using protocols previously described by us [22, 24, 41, 42, 52, 53]. Briefly, cell and EV lysates comprising 50 μg total protein were mixed with 4× Laemmli buffer and distilled water. This mixture was heated at 95 °C for 5 min using a heating block (Thermo Scientific). The samples and the premixed molecular weight markers (ladder, 250 kD-10 kD) were segregated on a 10% Mini-PROTEAN TGX Precast Protein Gels (Bio-Rad Laboratories, Inc.) at 120 V for 90 min using a PowerPac Basic setup (Bio-Rad Laboratories, Inc.). The proteins were transferred onto a 0.45 μm PVDF membrane using a transfer assembly (Sigma Aldrich) at 75 V and 300 mA for 90 min. Next, the membrane was washed with the 0.1%-Tween 20 Tris Buffered-saline (**T-TBS**) and was blocked using Odyssey blocking solution (Odyssey blocking buffer: T-TBS, 1:1) solution for an hour. The membrane was incubated overnight with either rabbit anti-Arf6 (1 mg/mL) at (1/1000 dilution) or rabbit anti-CD9 (0.2 µg/mL) primary antibody prepared in Odyssey blocking solution at 4°C. The membrane was again washed with T-TBS and incubated with anti-rabbit AF790 (0.05 μg/mL) in Odyssey blocking solution at room temperature for an hour. The washed membrane was scanned at 800 nm near-infrared channel using an Odyssey M imager (LI-COR Inc., Lincoln, NE).

### 2.10. Isolation of Mitotracker red (MitoT-EVs) labeled mitochondria containing EVs

Mitotracker deep red is a carbocyanine-based dye that stains mitochondria retaining the mitochondrial membrane potential [54, 55]. We pre-labeled the mitochondria of donor cells (NSC-34s or *d*NSC-34s) and then isolated MitoT-red labeled mitochondria containing EVs (refered as MitoT-lEVs and MitoT-sEVs). To isolate Mito-T labeled *d*NSC-34/NSC-34 EVs, confluent *d*NSC-34/NSC-34 cells were incubated with 100 nM Mitotracker deep red diluted in a neurobasal differentiation medium (to isolate *d*NSC-34-MitoT-EVs) or serum free DMEM high glucose medium ( to isolate NSC-34 MitoT-EVs) for 60 minutes. Post incubation, the cells were washed with PBS and were supplemented with fresh neurobasal differentiation medium or DMEM complete growth medium to isolate *d*NSC-34-MitoT-EVs and NSC-34 MitoT-EVs respectively, and incubated for 48 hours in a humidified incubator. Post-incubation, the EVs were isolated as per the differential ultracentrifugation protocol as described in the EV isolation method in section 2.4.

### 2.11. Flow cytometry analysis to quantify the uptake of MitoT-EV into the recipient NSC-34 or *d*NSC-34 cells

MitoT-lEVs and MitoT-sEVs derived from NSC-34 and *d*NSC-34 were isolated from the conditioned medium of either NSC-34 or *d*NSC-34 as described in section 2.4. NSC-34 cells were cultured in 24-well plates at 90,000 cells/well in a complete growth medium. Post-confluency, the NSC-34 cells were either differentiated to *d*NSC-34 or kept and treated with either MitoT-lEVs or MitoT-sEVs at 50, 75, and 100 µg EV protein/well in complete growth medium for 48 h in a humidified incubator. Untreated cells were used as gating control to exclude autofluorescence of cells. Post-treatment, the cells were washed with PBS followed by dissociation using TrypLE Express. The cells were suspended in 5% FBS + 2mM ethylenediaminetetraacetic acid (**EDTA)** in PBS, and collected in eppendorf tubes. For each sample, a 200 µL aliquot of the cell suspension was analyzed using an Attune NxT flow cytometer, and 30,000 events were recorded in FSC *vs.* SSC plots. MitoT-red fluorescence intensity was detected at an excitation/ emission of 644/665 nm and percentage signal intensities were recorded for data analysis as MitoT-red+ cells.

### 2.12. Fluorescence microscopy to detect EV-mediated mitochondrial transfer in recipient cells (NSC-34/*d*NSC-34)

MitoT-lEVs and MitoT-sEVs were isolated from NSC-34 cells’ conditioned medium following the previously mentioned EV isolation method in section 2.4. NSC-34 cells were cultured in a 96-well plate at 16,500 cells/well seeding density. At 80-85% confluency, cells were treated with MitoT-lEVs and MitoT-sEVs at 100 and 200 µg EV protein/well in the complete growth medium followed by 72 hours of incubation. Untreated NSC-34 cells were used as a control. Post-incubation, the cells were washed with PBS and incubated with a phenol red-free growth medium. Finally, cells were observed under an Olympus IX 73 epifluorescent inverted microscope (Olympus, Pittsburgh, PA) at an excitation/ emission setting of 644/665 nm and bright-field channel at 20× magnification. From each control and treatment group, at least three images were acquired and the total sum of grayscale signal intensities in the Cy5 channel was estimated using the Olympus CellSens software. The measured intensities were normalized with those of the untreated cells.

### 2.13. MTT assay to determine the effects of exposure of *d*NSC-34 *vs*. NSC-34 lEVs and *d*NSC-34 *vs.* NSC-34 sEVs to heat-stressed neurons

NSC-34 cells were seeded in a 96-well plate at 16,500 cells/well, and post confluency, the medium was switched to a Neurobasal differentiation medium. On day 3 of differentiation, *d*NSC-34 cells were exposed to Neurobasal differentiation medium pre-heated at 50.8 C for 30 minutes followed additional 30 minutes incubation at 50.8 C. The medium was removed and then replaced with 75 µg or 100 µg of *d*NSC-34 or NSC-34 lEVs or *d*NSC-34 or NSC-34 sEVs protein/well and incubated for 4 hours. Untreated, healthy, and untreated heat stress-exposed cells were used as controls. Post-incubation, the EV treatment was removed, and the heat-stressed neurons were supplemented with 100µL of fresh growth medium and 10µL of 12 mM MTT stock per well followed by 2 hours of incubation in a humidified incubator at 37 C. Next, 85 µL of neurobasal differentiation medium and MTT stock mixture was removed from each well followed by the addition of 50 µL of DMSO per well and incubated for 10 min at 37 C.

Finally, sample absorbance was measured at 570 nm wavelength using a SYNERGY HTX multi-mode reader (BioTek Instruments, Winooski, VT), and percent cell viability was calculated based on the following equation:

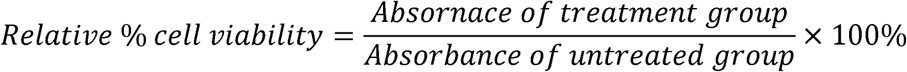

### 2.14. Seahorse assay to determine the effect of *d*NSC-34-derived lEVs and sEVs on heat-induced oxidative stress

NSC-34 cells were seeded in a seahorse XF96 well plate at 20,000 cells/well seeding density. Post-confluency, the cells were switched to neurobasal differentiation medium and incubated for 72 hours. At day 3 of differentiation, *d*NSC-34 refered to as neuron exposed to the neurobasal differentiation medium pre-heated at 50.8 C for 30 minutes followed by an additional 30 minute incubation at 50.8 C. The medium was replaced with 25 µg or 35 µg of lEVs or sEVs protein/well equivalent to 75 µg or 100 µg of lEVs or sEVs protein/well/in a 96-well plate. and incubated for 4 hours. The growth area of each well in a Seahorse plate is 0.114 cm^2^ and the treatment amount in each well was adjusted accordingly. Untreated heat stressed neurons were used as controls. Post-incubation, the treatment was replaced with 180 µL of pre-warmed DMEM-high glucose medium to perform the seahorse assay. After the measurement of baseline oxygen consumption rate (Basal **OCR**), 2.5 μmol/L oligomycin A and 0.7 μmol/L carbonyl cyanide-p-tri-fluoromethoxyphenyl-hydrazone were consecutively added to measure the proton leak and maximal OCR, respectively. The total protein content of the cells in each well was measured using Pierce BCA assay and Seahorse results were normalized to total cellular protein.

### 2.15. *In vivo* pilot study to detect MitoT-lEV-mitochondria in C57BL/6 mice

Young (2–3-month-old) male and female C57BL/6 mice were housed in the pathogen-free Animal Research Program (ARP) at Wake Forest University School of Medicine (WFUSM) under controlled conditions (20–23°C; 12:12-hour light-dark cycle). Transgenic ChAT-GFP mice, which express GFP in cholinergic neurons (ChAT[BAC]-eGFP, strain 007902) was purchased from Jackson Laboratory, Bar Harbor, Maine. Mice were fed standard chow ad libitum and had continuous access to drinking water. Muscle injections were performed under deep isoflurane anesthesia. All procedures complied with National Institutes of Health guidelines for the care and use of laboratory animals, and every effort was made to minimize suffering. The study was approved by the WFUSM Institutional Animal Care and Use Committee under protocol A23-012. To label mitochondria, confluent b.End3 BECs (90–95%) were incubated with 100 nM Mitotracker Deep Red in serum-free medium for 30 minutes. After incubation, cells were washed with PBS and cultured in fresh serum-free medium for 48 hours. MitoT-lEVs were then isolated using differential ultracentrifugation, as described in **section 2.4**. Following isolation, MitoT-lEVs were characterized using micro BCA for determining total EV protein content, DLS, and NTA were used to determine particle diameter, dispersity index, zeta potential, and EV particle concentration.We then injected the tibialis anterior (**TA**) and gastrocnemius (**GA**) muscles of C57BL/6 mice with increasing doses of MitoT-lEVs (1 × 10, 1 × 10, and 1 × 10 particles resuspended in a 5 µL injection volume), using three mice per dose group. Five days post-injection, spinal cord and muscle tissues were collected to evaluate whether MitoT-lEVs had been transported to the spinal cord, and whether any material remained at the neuromuscular junctions (**NMJs**) or the injection sites. Mice underwent cardiac perfusion for tissue fixation, followed by post-fixation, cryopreservation, sectioning, mounting, and imaging using differential interference contrast (**DIC**) and epifluorescence microscopy.

### 2.16. Statistical analysis

Statistical differences between the control groups and treatment groups or within the treatment groups were measured using one-way or two-way analysis of variance (ANOVA) at 95% confidence intervals using GraphPad Prism 10 (GraphPad Software, LLC). Tukey’s multiple comparison was performed for comparative analysis using one way and two way ANOVA. The different significance levels are indicated as follows: *p<0.05, **p<0.01, ***p<0.001 ****p<0.0001.

## 3. Results and discussion

### 3.1. The rationale for choosing NSC-34 cells

Neurons derived from the spinal cord have limited potential to grow and propagate. Therefore, under *in vitro* conditions, investigation of the neurotoxin-induced stress in motor neurons and studying their responses toward a therapeutic approach is challenging [56]. In 1992, Cashman and colleagues developed a series of neuroblastoma/spinal cord (**NSC-34**) hybrid cell lines and reported that the NSC-34 cell line resembled motor-neuron-like cells [57]. They reported that NSC-34 cells generated action potentials, and demonstrated synthesis, storage, and release of acetylcholine. The authors also reported that NSC-34 cells induced acetylcholine receptor clusters on cocultured myotubes, similar to those observed in neuronal development, suggesting that these cells may model the aspects of early neuromuscular synapse formation [57]. This cell line can be differentiated *in vitro* and used to study the pathophysiology of motor neurons in diseases such as ALS. In this study, we used NSC-34 cells as an *in vitro* ALS cell model and differentiated these cells into motor neurons using the protocol adapted from Madison *et al.* [51].

### 3.2. Differentiation of NSC-34 cells into neurons (*d*NSC-34/neurons)

We observed morphological changes in NSC-34 cells after switching them to Neurobasal differentiation medium. (**Fig.1a**). Post-differentiation, we observed the cells for five consecutive days under a phase microscope to study their morphological changes. We observed prominent neurite outgrowths (**Fig.1a**) consistent with the observations of Madison *et al.* [51]. Additionally, NSC-34 and *d*NSC-34 cells were stained with beta-III tubulin antibody, a neuronal marker to image neurite processes (**Fig.1b**).

**Fig. 1.**
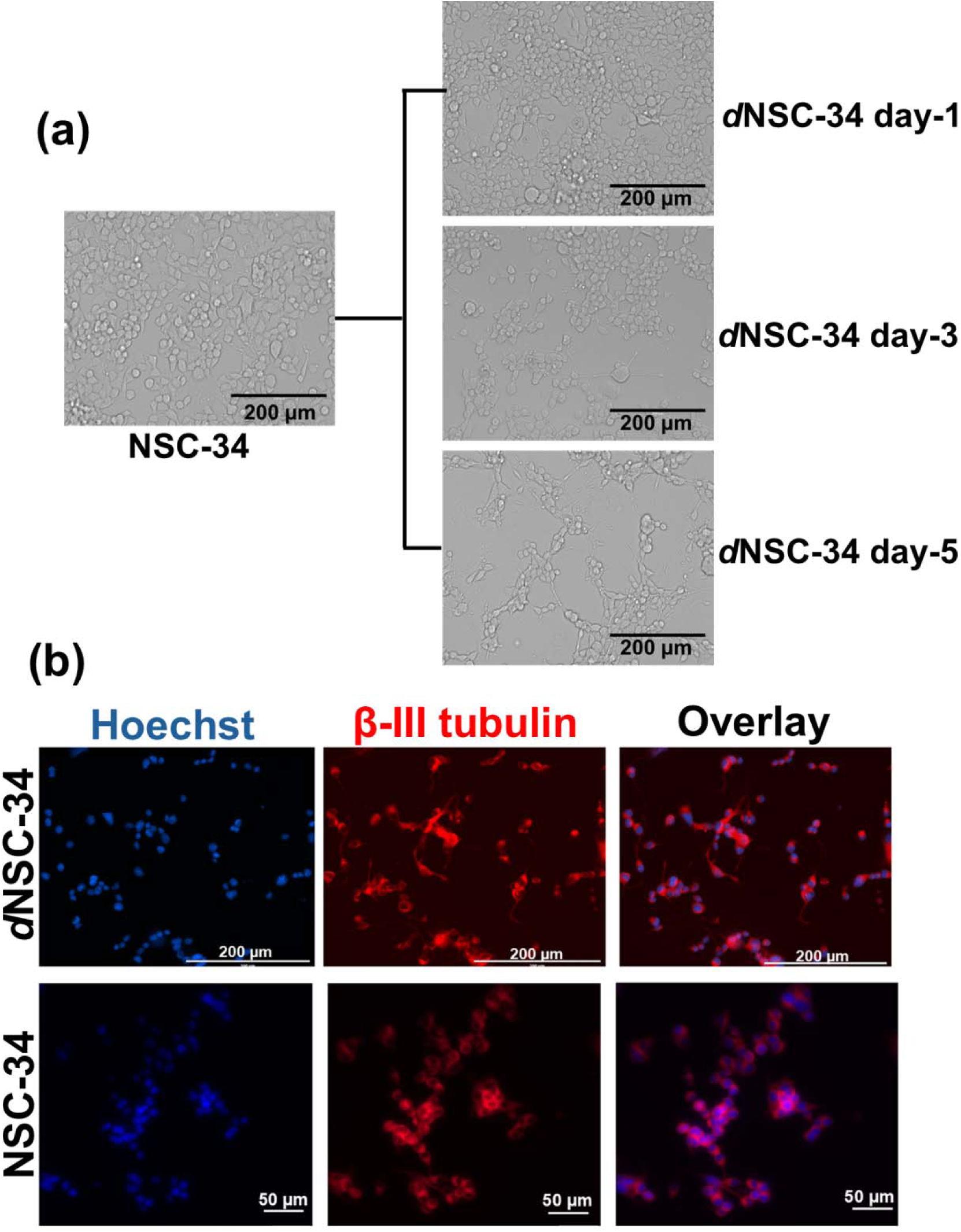
Microscopic analysis of NSC-34 vs. *d*NSC-34 cell morphology. **(a)** Microscopic image of basal NSC-34 cells showing no neurite outgrowths. NSC-34 cells were then shifted to Neurobasal differentiation medium and microscopic images were obtained for 5 consecutive days. Microscopic images, obtained on days 1,3 and 5 after differentiation demonstrate neurite outgrowths in *d*NSC-34 cells. (Scale bar: 200 µm) (**b)** NSC-34 and *d*NSC-34 cells were stained with beta-III tubulin antibody (neuronal marker) at 1:500 dilution and Hoechst dye at 10 µg/mL concentration (nuclear stain) followed by imaging under the RFP and DAPI channel, respectively. EVOS microscope was used to observe phase contrast images of *d*NSC-34 cells and Olympus epifluorescent microscope was used to image NSC-34 cells at 20*X* magnification. Scale bar for *d*NSC-34 cells: 200 µm and Scale bar for NSC-34 cells: 50 µm.

Class-III beta-tubulin was prominently expressed in the neurites of *d*NSC-34s **(Fig.1b)**. Microtubules are composed of alpha and beta-tubulin proteins [58] and out of the several existing isotypes of beta-tubulin, class III beta-tubulin has been considered a neuron-specific marker molecule [59]. Although beta-III tubulin was expressed in NSC-34 cells, neurite outgrowth was not observed **(Fig.1b)**. Overall, the above observations suggested that *d*NSC-34 demonstrated neuron-like characteristics. Therefore, this cell model was used for further *in vitro* experiments in this study.

### 3.3. Physicochemical characterization of NSC-34-derived lEVs and sEVs

Physicochemical characteristics of EVs govern their colloidal behavior and uptake into recipient cells. We used dynamic light scattering to measure the particle diameter, dispersity index, and zeta potential of EVs isolated from NSC-34 and *d*NSC-34 cells. Particle concentrations of EVs were measured using NTA. sEVs isolated from *d*NSC-34 and NSC-34 cells exhibited average particle diameters of 153.2 nm and 157.1 nm, respectively (**Fig.2a and Fig. S2a)**. lEVs isolated from *d*NSC-34s and NSC-34s showed average particle diameters of 217 ± 0.47 nm and 250.7 ± 3.0 nm, respectively (**Fig.2a and Fig. S2a**). This particle size data aligns with the published literature [32, 60–62] and minimal information for studies of extracellular vesicles (**MISEV 2023**) guidelines [32]. According to MISEV guidelines, EVs are a heterogeneous group of particles with sEVs ranging from 50-200 nm and lEVs ranging from 100-1000 nm. sEVs derived from *d*NSC-34 and NSC-34 cells showed anionic zeta potentials of - 25.9 mV and -32.9 mV, respectively (**Fig. 2a and Fig. S2a**). lEVs derived from *d*NSC-34s and NSC-34s demonstrated anionic zeta potentials of -28.5 mV and -35.6 mV, respectively (**Fig. 2a and Fig. S2a**). Zeta potential values indicates the surface charge of EVs. Published studies have shown the presence of anionic phosphatidylserine on the EV membrane along with phosphatidylcholine, sphingomyelin and glycoproteins [63–65]. The average particle concentrations of *d*NSC-34 and NSC-34 sEVs were 3.2×10^10^ and 1.6×10^11^ particles/mL, respectively (**Fig.2a and Fig.S2a**). lEVs isolated from *d*NSC-34s and NSC-34s exhibited particle concentrations of 2.1×10^11^ and 3.4×10^11^ particles/mL, respectively (**Fig.2a and Fig. S2a)**. The intensity-weighted particle size distribution plots of NSC-34 and *d*NSC-34-derived sEVs and lEVs are presented in **Fig. S1a-S1d**. Overall, the particle characteristics of EVs derived from NSC-34 and *d*NSC-34 cells were comparable, suggesting that the differentiation of NSC-34 cells into neurons did not markedly change EV biogenesis process or the resulting characteristics of EVs. For clarity, we have only presented the EV characteristics from *d*NSC-34 cells (neuronal EVs) as all subsequent studies use only neuronal EVs unless stated otherwise.

**Fig. 2.**
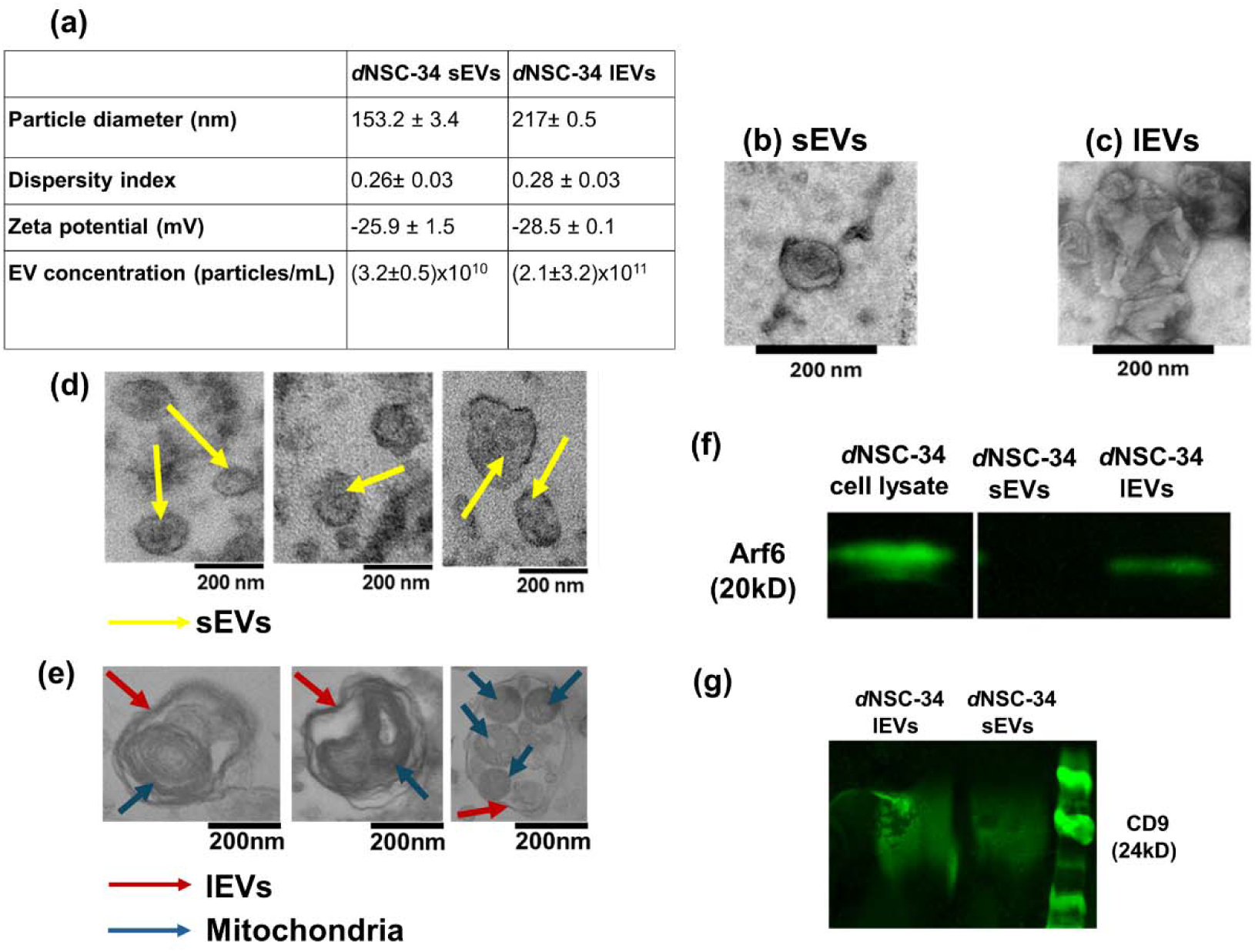
Physicochemical characterization, transmission electron microscopy, and western blot of *d*NSC-34 derived-sEVs and lEVs. **(a)** *d*NSC-34-derived sEVs and lEVs were characterized by measuring their particle diameter, dispersity indices, and zeta potential using a Malvern Zetasizer Pro DLS (Malvern Panalytical Inc., Westborough, PA). Samples were diluted at a 0.1 mg/ml concentration using either PBS or 10 mM HEPES buffer at pH 7.4 to measure particle size and zeta potential, respectively. Samples were run in triplicate and data were represented as mean± standard deviation (SD). Negative TEM images of *d*NSC-34-derived (**b)** sEVs and (**c)** lEVs. Representative TEM images of cross-sectioned (**d)** *d*NSC-34-sEVs and (**e)** *d*NSC-34-lEVs. lEVs (maroon arrow) contained one or mitochondria (electron-dense structure, blue arrows). sEVs (yellow arrows) showed no electron-dense structure in their lumen. (**f)**Western blotting to detect protein Arf6. Western blot analysis showed expression of Arf6 in *d*NSC-34 lEVs not in *d*NSC-34-sEVs. (**g**) Western blotting to detect CD9. Western blot analysis showed expression of CD9 in both *d*NSC-34 lEVs and sEVs. Raw blots are shown in supplementary **Fig. S5a and b**. For western blotting, the samples were electrophoresed on a sodium dodecyl sulfate-polyacrylamide gel followed by transfer to PVDF membrane. After treatment with blocking buffer, samples were incubated overnight with primary antibody at 4^0^C and with secondary antibody at room temperature. The blot was imaged using an Odyssey imager.

We observed the morphology of *d*NSC-34-EVs using TEM. Negative stain TEM images of lEVs showed larger particle sizes (**Fig. 2b**) (>200 nm) compared to sEVs (<200 nm) (**Fig. 2c)**. These findings were comparable to our previous study where brain endothelial cell-derived lEVs were larger than the sEVs [42]. Additionally, our findings align with several published literature [66–68]. S**upplementary Fig. S3a and S3b** are raw TEM images of negatively stained lEVs and sEVs. Arf6, a GTP-binding protein that contributes to the biogenesis and release of lEVs [69, 70] was expressed in lEVs but not sEVs suggesting the involvement of Arf6 in lEV biogenesis (**Fig. 2f and Fig. S2b**). Previously published literature also demonstrated the presence of Arf6 in lEVs derived from smooth muscle cells [71]. Furthermore, we determined the expression of CD9 marker protein in *d*NSC-34-EVs. Western blot data (**Fig. 2g and Supplementary Fig. S5b**) showed that CD9 was expressed in both sEVs and lEVs. CD9 is a member of tetraspannin family (group of plasma membrane proteins) considered as a EV specific biomarker because of its high enrichment on EV membranes [72, 73]. Our findings were comparable with several studies which demonstrated enrichment of EVs with CD9 [73–75]. It is important to note that although CD9 is a sEV specific biomarker [62, 72, 76], lEVs too have been reported to express CD9 [77, 78].

### 3.4. Presence of mitochondria in lEVs

Multiple cross-sectioned TEM images of *d*NSC-34-derived EVs were analyzed to study if lEVs from *d*NSC-34 cells entrap mitochondria similar to our prior reports of mitochondria in BEC-derived lEVs [22–24]. We observed that the lumen of lEVs (**marron arrows Fig.2e**) contained one or multiple electron-dense structures (**blue arrows Fig.2e**) suggesting the presence of mitochondria. However, these electron-dense structures were not detected in sEVs suggesting the lack of mitochondria (**yellow arrow Fig. 2d**). Uncropped cross-sectioned TEM images of sEVs and lEVs are shown in **supplementary Fig S4a and S4b**. This observation was comparable to our previous studies which found that BEC-derived lEVs, but not sEVs contained mitochondria [37, 39, 57]. In addition to our prior observations, our findings of mitochondria in lEVs also align with several published works where the author has shown the presence of mitochondria in lEVs derived from human endothelial progenitor cells, stem cell-derived cardiomyocytes, and mesenchymal stem cells [26, 35, 36, 79]. The observed morphology of mitochondria in the lEVs also aligned with the morphology reported by other groups [80–82]. It should be noted that the morphology of mitochondria in the TEM images can be impacted by the extensive sample processing and the high-speed centrifuging steps involved in sectioning EVs [83].

### 3.5. lEV-mediated mitochondrial transfer into recipient neurons

In the current study, we wanted to determine whether *d*NSC-34-derived lEVs can transfer their mitochondrial load to the recipient neurons using flow cytometry and fluorescence microscopy analysis. Mitotracker deep red (**MitoT**) is a carbocyanine-based dye that specifically stains mitochondria depending on mitochondrial membrane potential [54, 55]. We pre-labeled the mitochondria of donor cells (NSC-34 and *d*NSC-34) and then isolated MitoT-labeled mitochondria containing EVs (**MitoT-lEVs and MitoT-sEVs**). Then, we treated NSC-34s/*d*NSC-34s with MitoT-lEVs or -sEVs at doses ranging from 50 to 100µg MitoT-EV protein/well (**Figure 3a-c**). Previously we have demonstrated that lEVs transfer their mitochondrial content to the recipient human BECs or mouse BECs and improve their mitochondrial bioenergetics and cell survival under oxygen-glucose-deprived conditions [24, 41, 42, 52].

**Fig. 3.**
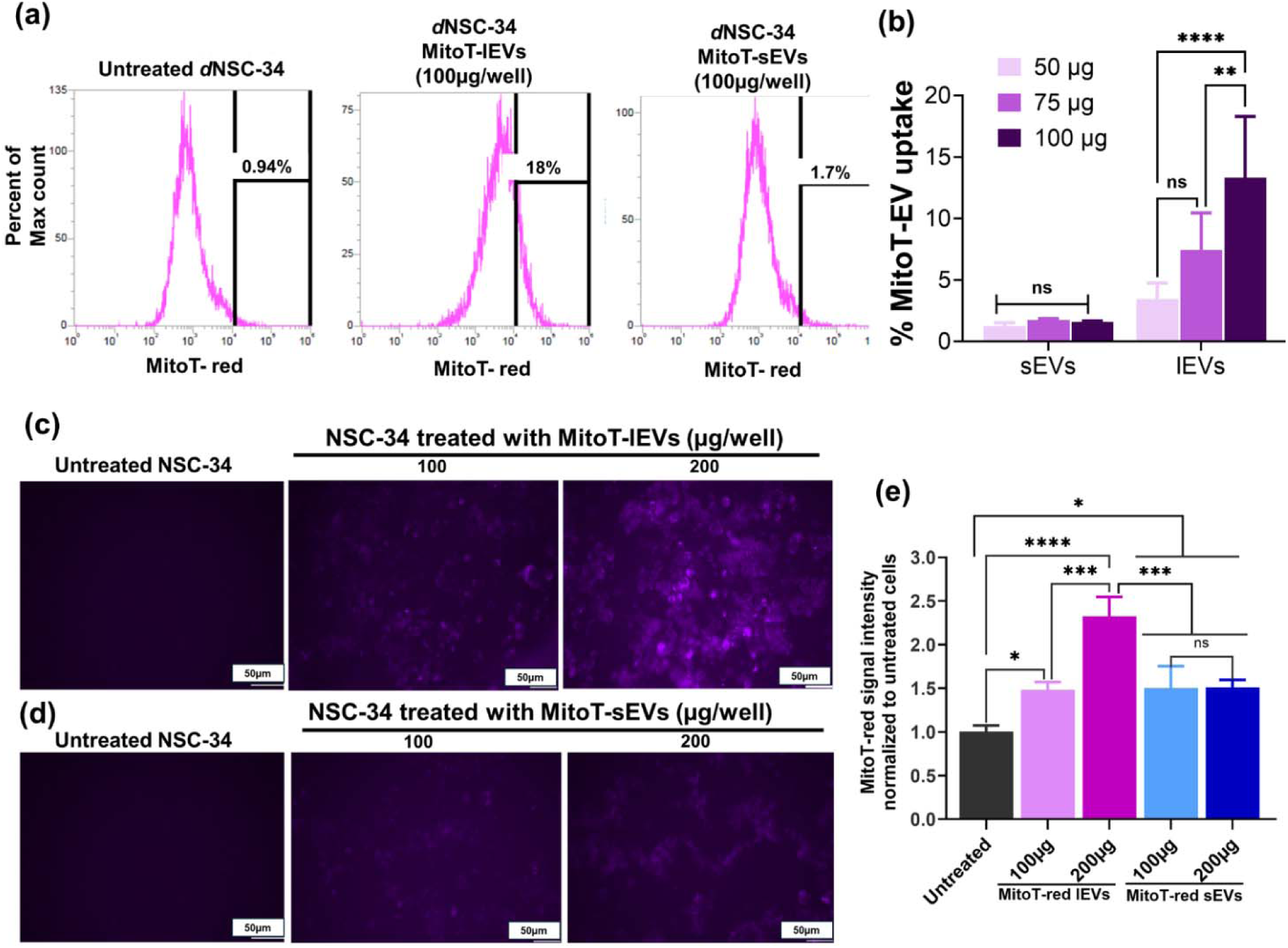
**EV-mediated mitochondrial transfer into recipient neurons** The representative histogram plots of *d*NSC-34 cells treated at dose of 100 µg/well of *d*NSC-34**-**MitoT-lEVs or **(b)** *d*NSC-34-MitoT-sEVs were collected using a 674/10-nm side scatter filter in the Attune flow cytometer. **(a)** Untreated *d*NSC-34 cells were used as controls to gate the background signals in histograms. As a representative image of mitochondrial transfer into recipient neurons, we reported the histogram plot of recipient *d*NSC-34 cells treated with 100µg/well of MitoT-lEVs and MitoT-sEVs derived from *d*NSC-34 cells (**b)** Quantification of EV-mediated mitochondrial transfer into recipient *d*NSC-34 cells was obtained from multiple histogram plots of recipient *d*NSC-34 cells treated with MitoT-lEVs or MitoT-sEVs. Data represented as mean ± SD of n=4. **(c) and (d)** Fluorescence microscopic analysis of EV-mediated mitochondrial transfer. Recipient NSC-34 cells were cultured in 96-well plate at 16,500 cells/well seeding density until they became 80-85% confluent. Subsequently, cells were treated with MitoT-sEVs or MitoT-lEVs at indicated doses for 72 hours. Post-incubation, MitoT-EV treated NSC-34 cells were imaged using Olympus fluorescence microscopy for detecting EV-mitochondria (purple puncta) (**c)** Recipient NSC-34 cells treated with MitoT-lEVs at indicated doses for 72 hours showed relatively higher MitoT-red fluorescence signal. (**d)** Recipient NSC-34 cells treated with MitoT-sEVs showed very faint fluorescence signal. Images were taken under Cy5 channel at 20X maginification. Scale bar: 50µm. (**e**) Quantification of EV mitochondria transfer into recipient NSC-34 cells. NSC-34 cells were treated with the indicated doses of MitoT-red lEVs or MitoT-red sEVs for 72 h. From each control and treatment group, at least three images were acquired and the total sum of grayscale signal intensities in the Cy5 channel was estimated using Olympus CellSens software. The measured intensities were normalized with those of the untreated cells.

Therefore, the recipient NSC-34 and *d*NSC-34 cells were treated with NSC-34/***d***NSC-34-derived MitoT-lEVs or MitoT-sEVs at 50 µg, 75 µg, and 100 µg EV protein/well in a 24-well plate for 48 h. We measured the percentage of cells positive for the MitoT-red fluorescence signal using flow cytometry. The recipient cells positive for MitoT-red fluorescence signal indicate the percentage of total cells internalizing MitoT-EVs and therefore the data are represented as %MitoT-EV uptake in (**Fig.3b and Fig. S6a-S6c)**. We analyzed the histogram plots of *d*NSC-34 cells treated at indicated doses of MitoT-lEVs and MitoT-sEVs derived from *d*NSC-34 (**Fig. 3a and 3b).** We observed a dose-dependent increase in MitoT-red positive cells after MitoT-lEV treatment compared to the levels noted with MitoT-sEV-treated cells suggesting the lEV-mediated mitochondrial transfer into recipient cells (**Fig.3b and Fig. S6a-S6c**). Around 3.4% of recipient *d*NSC-34 cells were positive for MitoT-red when treated with 50 µg of *d*NSC-34-derived MitoT-lEVs, whereas 7.4% of cells showed a MitoT-red positive signal at 75µg of MitoT-lEVs dose(**Fig.3b)**. The percentages of MitoT-red positive cells significantly (p<0.0001) increased to 13.3% at 100 µg of lEVs dose per well (**Fig.3b**). However, the fraction of MitoT-red positive *d*NSC-34 cells was significantly low when treated with *d*NSC-34-MitoT-sEVs ranging from 1.2% to 1.6% at the three tested doses (50 µg, 75 µg, and 100 µg of MitoT-red sEVs per well) (**Fig.3a and 3b**). This experiment was repeated with three other pairs of EVs and recipient cells: recipient NSC-34s treated with *d*NSC-34-derived EVs; recipient *d*NSC-34s treated with NSC-34-derived EVs and recipient NSC-34s treated with NSC-34-derived EVs (**Supplementary figure S6a-S6c**). The same trend of lEV-mediated mitochondrial transfer was observed in those recipient NSC-34/*d*NSC-34 cells (**Supplementary Fig.S6a-c)**. In contrast, sEVs did not demonstrate any substantial uptake by recipient NSC-34s/*d*NSC-34s as shown in (**Fig.S6a-S6c)**. These findings correlate with the cross-sectioned TEM images of EVs that demonstrated the presence of electron-dense mitochondria in lEV but not sEV lumen (**Fig. 2d and 2e**).

We performed fluorescence microscopy analysis further to investigate the lEV-mediated mitochondria transfer into recipient NSC-34 cells. In this study, recipient NSC-34 cells were incubated with MitoT-lEVs or MitoT-sEVs at a dose of 100 or 200 µg for 72 hours. Post-incubation, cells were imaged using the Olympus fluorescence microscopy under the Cy5 channel for detecting EV-mitochondria (purple puncta) (**Fig. 3c and 3d**). A faint signal was observed in recipient NSC-34 cells treated with 100 µg of MitoT-lEV protein (**Fig. 3c**). However, a higher level of MitoT-red signal was observed in recipient NSC-34 cells treated with 200 µg/well of MitoT-lEVs suggesting efficient mitochondrial transfer via lEVs at this does (**Fig. 3c**). In contrast, recipient NSC-34 treated with 100 or 200 µg/well of MitoT-sEVs protein for 72 hours, showed a very faint purple puncta suggesting lower mitochondrial load (**Fig. 3d**). The extent of EV mitochondria delivery into recipient NSC-34 cells was quantified based on the relative fluorescence intensity of Mitotracker Red dye (**Fig. 3e**). The data in **Figure 3e** demonstrated that recipient NSC-34 cells treated with 200 µg/well of MitoT-lEVs showed a significantly greater fluorescence intensity compared to 100 µg/well of lEVs suggesting a dose-dependent delivery of lEV mitochondria. In contrast, recipient cells treated with 100 µg/well and 200 µg/well of MitoT-sEVs showed significantly lower fluorescence intensities compared to cells exposed to MitoT-lEVs. These findings suggests that lEVs transfer their mitochondrial load to a greater extent than sEVs.

Here, we evaluated whether NSC-34/*d*NSC-34-EVs transfer their mitochondrial load to the recipient cells. Our data demonstrated that irrespective of the experimental pairs, lEV-treated recipient NSC-34/*d*NSC-34 cells showed significantly greater MitoT-red fluorescence intensity, while sEV-treated recipient cells showed lower levels of MitoT positive signals. These findings suggested that lEV but not sEV contains mitochondria and can transfer its mitochondrial load to the recipient NSC-34s/*d*NSC-34s. This aligns well with the existing literature on the ability of EVs to transfer their mitochondrial content into the treated cells [22, 41, 42, 52, 84]. Collectively, these data suggested that neuron-derived EVs can transfer their mitochondrial load into the recipient neurons.

### 3.6. Effect of EV treatment on heat-stressed recipient neurons

After confirming the presence of mitochondria in *d*NSC-34 lEVs and the subsequent transfer of lEV-mitochondria into the recipient neurons, we studied the functional effects of this lEV-mediated mitochondrial transfer in heat-stressed neurons. Although we did not find any trace of mitochondria in sEVs, we wanted to evaluate its functional effect. Neurons (*d*NSC-34 cells) were exposed to 50.8 °C heat to induce oxidative stress. Heat stress can impact the physiology of motor neurons by inducing oxidative stress, excessive intracellular calcium load, protein denaturation, and mitochondrial dysfunction that represent some pathophysiologic features of ALS [85]. Neurons exposed to heat stress showed morphological changes accompanied by a loss of neurite outgrowth **(Fig. S7c)**.

To further substantiate the potential of *d*NSC-34-derived EVs, we exposed heat-stressed neurons with NSC-34 or *d*NSC-34-derived EVs and measured the outcomes of EV exposure by performing a MTT assay (**Fig. 4**). MTT assay is a colorimetric assay that determines cell viability based on the activity of NAD(P)H-dependent cellular oxidoreductase enzymes present in the inner mitochondrial membrane and catalyze several reactions of the electron transport chain to produce ATP [86]. If the cells are viable, oxidoreductase will reduce the soluble MTT tetrazolium salt to form insoluble formazan. Finally, formazan is solubilized, and cell viability is determined by measuring the absorbance at a wavelength of 570 nm. Considering the correlation between cell viability and mitochondrial function, we performed the MTT assay to evaluate the effect of mitochondria-containing EVs on heat-stressed neurons. MTT assay results demonstrated that *d*NSC-34 cell-derived lEVs and sEVs showed significant improvement in cell viability in heat-stressed neurons compared to NSC-34 derived EVs. **Fig. 4** showed that the cell viability of control/untreated heat-stressed neurons that reduced to 32% significantly increased to 61% when treated with 75 µg of *d*NSC-34 lEVs/well. 100 µg of *d*NSC-34-lEV protein treatment showed a significant increase to 85% in cell viability. However, for 100 µg of NSC-34 lEVs we did not observe any significant increase in cell viability. Additionally, we observed a significant effect of 75 µg and 100 µg of *d*NSC-34 sEV treatment on the cell viability of heat stressed neurons. Neuron cell viability significantly increased to 65% and 83% when treated with 75 µg and 100 µg of *d*NSC-34 sEVs, respectively. However, doses of 75 µg and 100 µg of NSC-34 sEVs did not show any substantial improvement in cell viability. <u>Collectively, we observed that</u> *<u>d</u>*<u>NSC-34 cell derived lEVs and sEVs showed significant improvement in cell viability of the</u> <u>stressed neurons compared to NSC-34 derived lEVs and sEVs</u>. <u>These findings are important as</u> <u>they demonstrate that EVs from *d*NSC-34—a neuronal source provide superior protective effects</u> <u>compared to EVs from undifferentiated NSC-34 cells.</u> Interestingly, both *d*NSC-34-derived sEV and lEV treatments showed an increase in cell viability. sEVs have been reported to contain mitochondrial DNA and mitochondrial proteins [34, 36, 87, 88] that might contribute to increasing the cell viability in stressed neurons. Noteworthy, sEVs are reported to contain a cocktail of heat shock proteins (**HSPs**) in their lumen that may have an impact on improving cell viability upon cell duress [41].

**Fig. 4.**
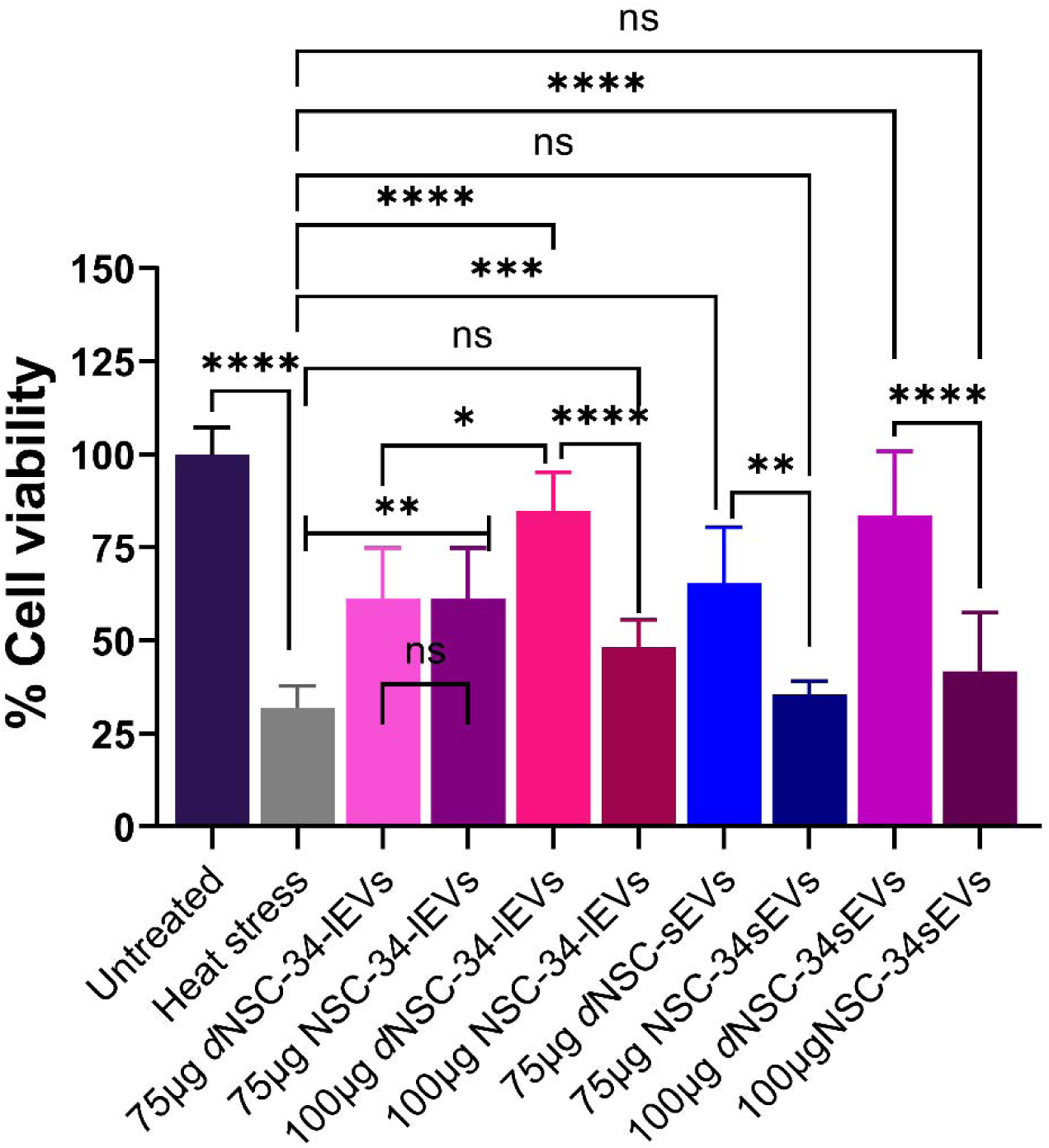
**MTT assay to determine the effect of *d*NSC-34 derived EVs on heat-stressed neurons** Neurons were cultured in a 96-well plate at a seeding density of 16,500 cells/well. Neurons were treated with a neurobasal differentiation medium pre-heated at 50.8lJC or left untreated for 30 minutes. Post-treatment, the heat stress condition was replaced with either *d*NSC-34 or NSC-34 derived lEVs or sEVs at 75 or 100 µg EV protein per well and incubated for 4 hours in a humidified incubator at 37°C. Finally, the effect of lEVs and sEVs on heat-stressed neurons was determined by performing a MTT assay. Data were represented as mean ± SD (n =6). Statistical analysis was performed by using one-way ANOVA followed by Tukey multiple comparison test. ****p<0.0001; **p<0.01 ^ns^non-significant.

Our previously published proteomic analysis of BEC-EVs revealed an enrichment of several endogenous HSPs: (HSP70, HSP71A, HSPA8, HSPA5, HSP105, HSP90A/B and HSPβ1) [21]. Guescini *et al*. found that sEVs derived from astrocytes and glioblastoma cells carry mitochondrial DNA contributing to inter-cellular communication and with potential implications for several neurodegenerative diseases [34]. Guescini *et al*. also reported that C2C12 myoblast also releases sEVs carrying mitochondrial DNA and mitochondrial proteins influencing several transduction mechanisms in the recipient cells [87]. Besides, sEVs are reported to contain different HSPs [89, 90] in their lumen which might have an impact on improving cell viability under cell duress. Shi *et al*. characterized sEVs derived from THP-1 cells. This study reported the presence of HSP27 in sEVs that stimulated NF-κB and release of IL-10. These data demonstrated the role of HSP27 in initiating anti-inflammatory effects [90]. Based on these published reports, we presume that sEVs derived from *d*NSC-34 cells may carry mitochondrial DNA, mitochondrial proteins as well as a cocktail of HSPs that are likely involved in the observed protective effects.

To further understand the effects of EV exposure on the resulting bioenergetics of EV-treated neurons, we performed a Seahorse assay. Seahorse extracellular flux assay is a state-of-the-art technique to measure mitochondrial function in cells and biological fluids via measurement of oxygen consumption rate (**OCR**) and extracellular acidification rate (**ECAR**) [91–95]. OCR values reflect mitochondrial respiration and ECAR values are reflective of glycolytic capacity [96]. Intriguingly, **Fig. 5a** showed that recipient neurons treated with 25 and 35 µg doses of both lEVs and sEVs significantly increased basal OCR and maximal OCR. The magnitude of OCR increases in the case of sEVs seemed to be greater than those observed in lEV-treated cells though the differences were not always statistically significant. However, the proton leak was significantly lower in recipient neurons treated with both doses of lEVs compared to sEVs. This suggests that the OCR increases observed with lEV treatment but not sEVs is coupled to oxidative phosphorylation.

**Fig. 5.**
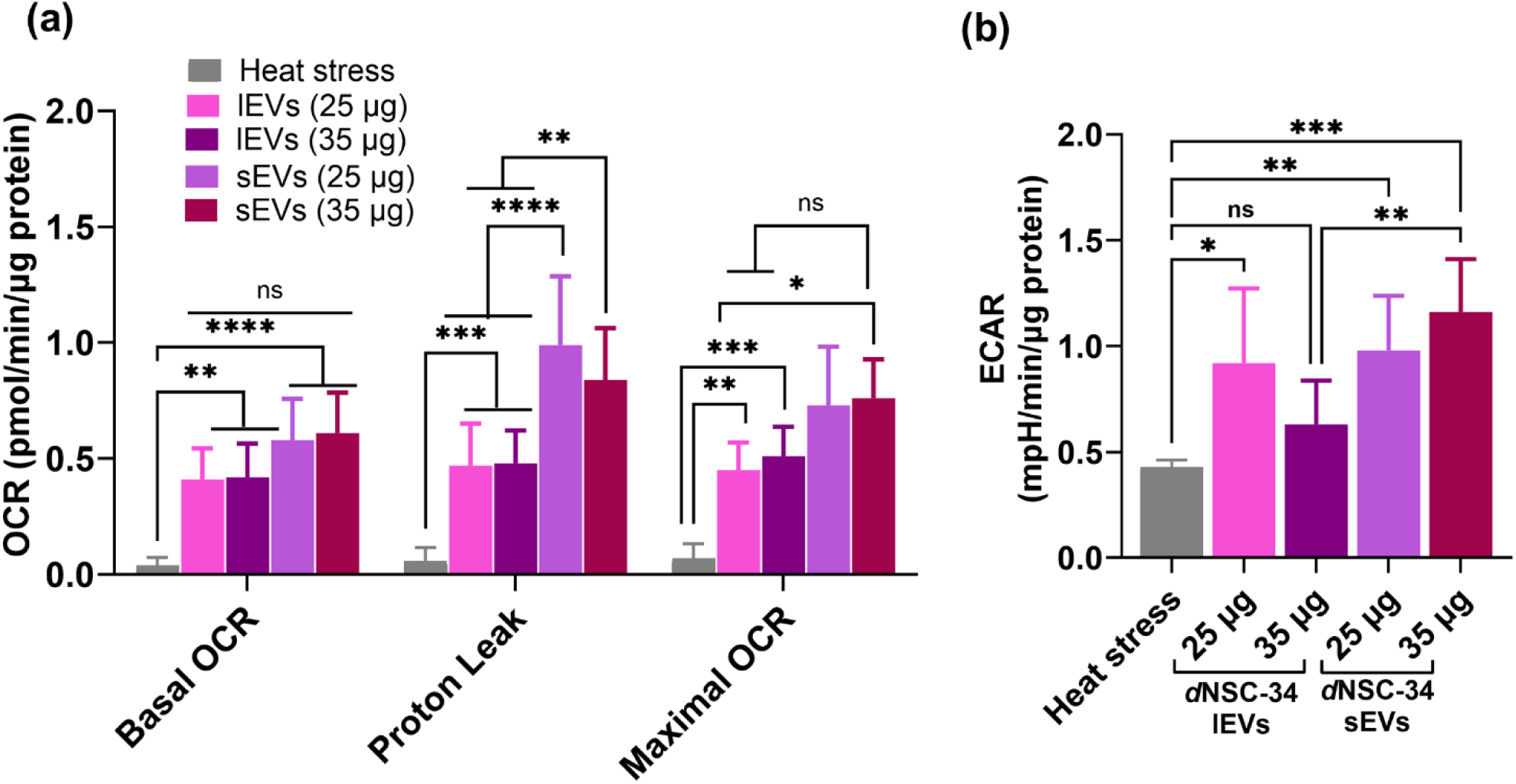
Seahorse assay to determine the effect of *d*NSC-34-derived EVs on heat-stressed neurons. *d*NSC-34 cells were seeded in a Seahorse XF96 well plate at 20,000 cells/well seeding density. Post-confluency, the cells were switched to Neurobasal differentiation medium and incubated for 72 hours. At day 3 of differentiation, *d*NSC-34 cells were exposed to heat stress followed by treatment with lEVs and sEVs at indicated doses for four hours. Post-treatment, a seahorse assay was performed. Subsequently, **(a)** basal oxygen consumption rate (basal OCR), proton leak and maximum OCR and **(b)** basal glycolytic rate was determined by measuring extracellular acidification rate (ECAR) were analyzed using the Seahorse XFe96 analyzer. Data represents mean ± SD (n = 6). * p < 0.05, ** p < 0.01, *** p < 0.001, **** p < 0.0001.

According to the working principle of the Seahorse assay, proton leak is determined after exposing the cells to oligomycin: an ATP synthase inhibitor. ATP synthase utilizes the proton gradient produced by the electron transport chain to generate ATP. During this ATP production, ATP synthase pumps out the proton from the mitochondria’s intermembrane space to the mitochondrial matrix, thereby maintaining mitochondrial membrane potential. When ATP synthase is blocked, it fails to maintain the proton gradient across the mitochondrial membrane. However, under certain circumstances, protons (for example, in the event of damaged mitochondrial membranes) can flow back to the mitochondrial matrix without passing through ATP synthase. This proton leak contributes to OCR values not associated with ATP synthesis. Based on this working principle, we expected that if lEVs transferred their mitochondria to the recipient neurons, the proton leak may be significantly lower compared to neurons treated with sEVs. As expected, neurons treated with sEVs showed a greater magnitude of proton leak compared to lEV-treated cells. We also analyzed the effect of EV exposure on recipient cell glycolysis based on basal ECAR. We observed an increase in basal glycolytic rate in neurons treated with 75 µg of lEVs whereas 100 µg of *d*NSC-34-lEVs did not show any significant increase in basal ECAR. In contrast, for both doses of *d*NSC-34-sEVs, we observed a significant increase in basal glycolytic rates. Higher ECAR values in sEV-treated neurons suggested that the energy production of neurons treated with sEVs was mostly dependent on glycolysis instead of oxidative phosphorylation. In contrast, heat-stressed neurons treated with 35 µg of lEVs showed a reduction in basal ECAR value suggesting that the cells were able to produce energy through oxidative phosphorylation via effects of the lEV mitochondrial load. However, under stressed conditions, cells likely produce energy for their survival using different pathways. This may explain the overall increase in ECAR value for all EV-treatment groups.

Collectively, Seahorse data showed that both subtypes of EVs increase the mitochondrial function of heat-stressed neurons. lEVs likely improved the mitochondrial respiration of heat-stressed neurons via the transfer of innate lEV mitochondria. In contrast, although sEVs do not contain mitochondria, they significantly increased the mitochondrial respiration of stressed neurons albeit at the expense of a *greater proton leak* compared to cells treated with lEVs. Though sEVs do not carry mitochondria [42, 72, 97], sEVs are reported to carry mitochondrial components such as mitochondrial DNA and mitochondrial proteins [34, 98]. Noteworthy, sEVs are also reported to contain different types of HSPs in their lumen that may have an impact on improving mitochondrial respiration under cell duress [41]. Our further studies will complete a proteomic profiling of both subtypes of EVs to determine the differences in their molecular and mitochondrial cargo.

EVs are known to contain mitochondria, mitochondrial DNA, and proteins when they are released from the cells [34, 85, 87, 88]. Numerous studies have demonstrated that EVs can transfer their mitochondrial cargo load into the recipient cells in different conditions. For instance, adipose-derived mesenchymal stem cell-derived exosomes transferred their mitochondrial load to lipopolysaccharide (**LPS**)-stimulated alveolar macrophages and increased OXPHOS activity, mitochondrial membrane potential, and ATP generation [99]. Additionally, in a mice model of LPS-induced acute lung injury, the authors demonstrated the anti-inflammatory and tissue-protective role of mitochondria in adipose-derived mesenchymal stem cell-derived exosomes [99]. In another study, EVs isolated from human-induced pluripotent stem cell-derived cardiomyocytes improved mitochondrial bioenergetics in the recipient hypoxic human-induced pluripotent stem cell-derived cardiomyocytes *in vitro* and in an *in vivo* murine myocardial infarction model. They demonstrated that the EV mitochondria were transferred into the recipient cells and fused with the endogenous mitochondrial network, which was likely responsible for the functional effects [35]. Our observation also aligns with these research findings. Therefore, the data collectively suggests that lEVs and sEVs can enhance the viability of heat-stressed neurons by transferring their mitochondrial components.

### 3.7. Efficient dose-dependent uptake of MitoT-lEVs by mouse motor neurons *in vivo* upon intramuscular injection

We completed a pilot study in healthy C57BL/6 mice to determine if mitochondria-containing large EVs can be transported to mouse motor neurons in the spinal cord. In this experiment, we used MitoT-labeled large EVs derived from mouse brain endothelial cells (**BECs**) based on the rationale that BEC-derived EVs were safely tolerated upon administration in C57BL/6 mice in our prior studies [24]. C57BL/6 mice were intramuscularly injected with MitoT-labeled lEVs in the right tibialis anterior (**TA**) and gastrocnemius (**GA**) muscles at three different doses of MitoT-labeled EVs: 1×10^6^, 1×10^7^ and 1×10^8^ EV particles. Mice injected with the highest dose of MitoT-lEVs (1 × 10 EVs) resulted in successful delivery of labeled mitochondria to the lumbar spinal cord. Numerous spinal cord neurons displayed the characteristic fluorescent puncta in the motor neuron regions (circled) as well as beyond (**Fig. 6b** and **6d**). Fluorescence was also clearly observed on the contralateral side of the injection (**Fig. 6c**), and extensive labeling of dorsal neurons was noted. At the injection sites in both TA and GA muscles, MitoT-lEVs had almost completely cleared by the time of tissue collection. Injections of medium and low doses of MitoT-lEVs in three mice per group also resulted in spinal cord labeling. As shown in **Fig. 6f**, the medium dose produced more robust neuronal labeling compared to the low dose (**Fig. 6h**), where the signal was faint and difficult to distinguish from background. To confirm motor neuron-specific uptake, we utilized transgenic ChAT-GFP mice, which express GFP in cholinergic neurons. **Fig. 6e–g** corresponds to a ChAT-GFP mouse injected with the medium dose of MitoT-lEVs. **Fig. 6e** shows the GFP-labeled motor neurons in the ventral horn, while **Fig. 6f** demonstrates strong MitoT-lEV signal in the same region, indicating successful delivery of mitochondria to spinal motor neurons. The overlay image (**Fig. 6g**) confirms colocalization of the MitoTracker Deep Red signal with ChAT-GFP-positive motor neurons, suggesting specific uptake of MitoT-lEVs by these cells. In contrast, the low dose resulted in only faint labeling of motor neurons (**Fig. 6h**), indicating a reduced extent of mitochondrial transfer at this dose. We interpret that MitoTracker-labeled lEVs are retrogradely taken up at the neuromuscular junctions and muscle spindle—structures innervated by group Ia, group II afferents, and gamma motor neurons—resulting in labeling of both large and small motor neurons, as well as sensory neurons in the spinal cord. Based on these findings, we conclude that spinal motor neurons are capable of internalizing MitoT-lEVs-mitochondria. Furthermore, this study demonstrated that medium-dose administration provides effective neuronal labeling without any evident advantage from the higher dose. Therefore, we propose to use the medium dose of MitoT-lEVs for subsequent experiments. In future studies, we will apply this approach to the SOD1^G93A^ transgenic mouse model of ALS to evaluate the therapeutic efficacy of lEV-delivered mitochondria.

**Fig. 6.**
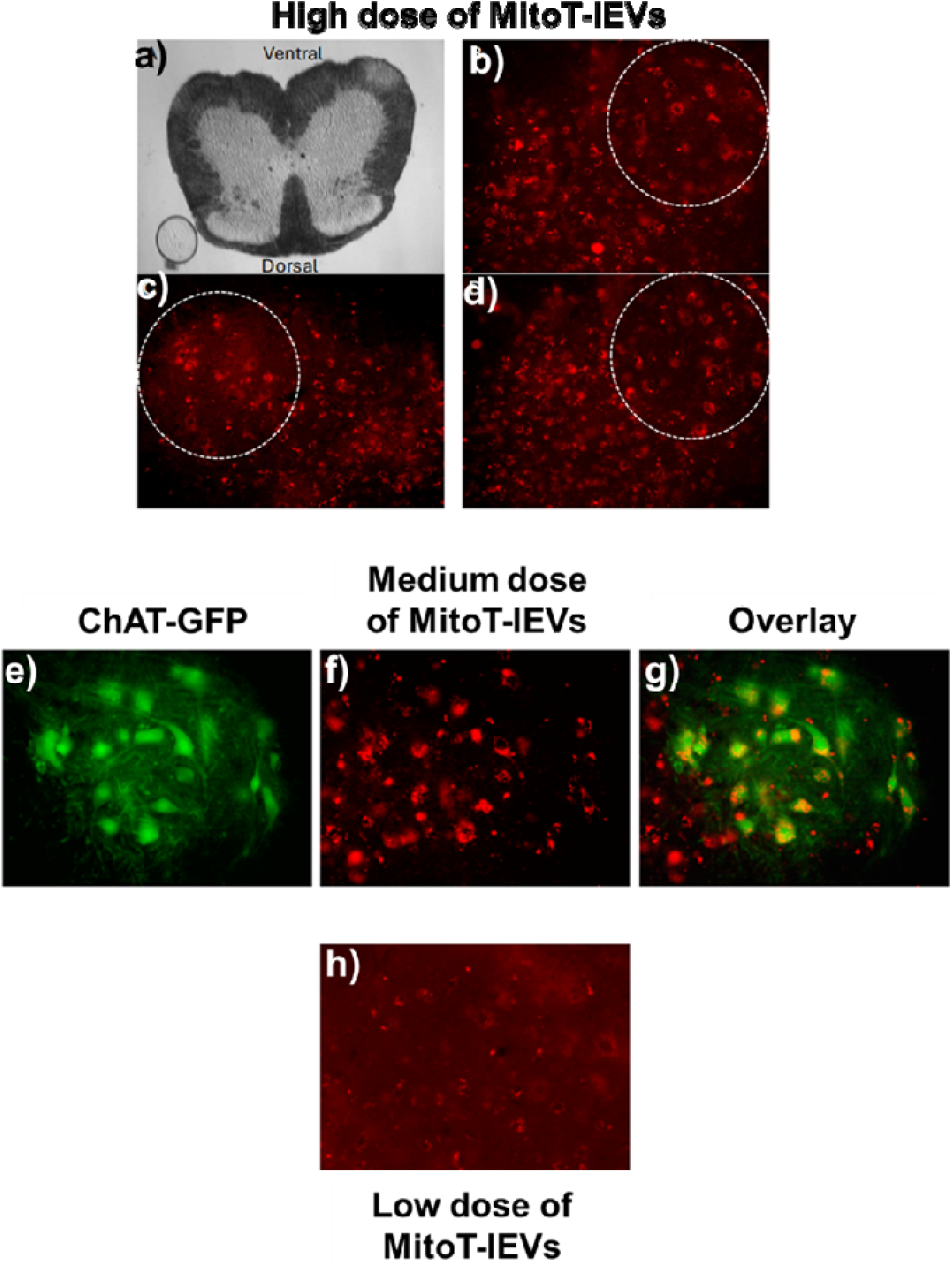
***In vivo* pilot study to detect MitoT-lEV mitochondria in C57BL6 mice** MitoT-lEVs were isolated from b.End3 BECs. The tibialis anterior and gastrocnemius muscles of C57BL/6 mice were then injected with three different doses of MitoT-lEVs (1 × 10LJ, 1 × 10LJ, and 1 × 10LJ particles per 5 µL injection; *n* = 3 mice per dose). **(a)** Dorsal and ventral spinal cord images were captured using standard differential interference contrast optics. (**b–d**) Epifluorescence images of lumbar spinal cord sections from mice injected with the high dose (1 × 10LJ). Numerous spinal neurons show characteristic fluorescent puncta in the motor neuron region (circled) and beyond (b, d), including the contralateral side of injection (c). (**e–g**) Epifluorescence images of lumbar spinal cord from a ChAT-GFP transgenic mouse injected with the medium dose (1 × 10lJ). (**e**) Transgenically labeled ChAT-positive motor neurons in the ventral horn. (**f**) MitoT-lEV signal alone, highlighting uptake in the same region. (**g**) Overlay of (e) and (f) shows that while MitoT-lEVs are present in various neurons, they are specifically taken up by motor neurons. (**h**) Lumbar spinal cord section from a mouse injected with the low dose (1 × 10lJ), showing faint MitoT-Red signal. All fluorescence images are 0.698 mm × 0.532 mm and were acquired using Cy5 excitation/emission filters at 20× magnification with a resolution of 0.519 µm/pixel. Exposure time was consistent across all mice and doses (500 ms).

## 4. Limitations of the study

In this study, we have found that both neuronal lEVs and sEVs can increase mitochondrial function albeit concomitant with a greater proton leak in the sEV-treated cells—in contrast to our prior findings where only BEC-derived lEVs but not sEVs showed increased mitochondrial function in the recipient BECs [21, 23, 24]. The observed functional effects (increased mitochondrial function, glycolytic capacity and overall cell survival—**Figures 4 and 5**) in sEV-treated cells is certainly interesting but these findings warrant additional investigation into the molecular cargo of sEVs *vs*. lEVs. sEVs did not show presence of mitochondria (**Figure 2**), but it is likely that mitochondrial components such as mitochondrial DNA (**mtDNA**) and mitochondrial proteins can result in survival-promoting effects in the recipient neurons. In fact, other studies have reported that sEVs contain mtDNA and mitochondrial proteins [34, 36, 87, 90]. We are currently optimizing protocols to quantify mtDNA in EVs using qPCR and experiments to compare the proteomic profiles of neuronal sEVs *vs*. lEVs are underway in our laboratory.

## 5. Conclusions

We have developed and studied the physicochemical characteristics of neuronal EVs derived from differentiated NSC-34 cells. For the first time, we have demonstrated increased mitochondrial function in the recipient neurons treated with either neuronal sEVs or lEVs— suggesting that mitochondrial cargo in EVs can exert bioenergetic responses. We anticipate that further studies on defining the differences between molecular cargo among neuronal sEVs *vs*. lEVs will shed light on the mechanism behind these bioenergetic responses. The promising results from a pilot *in vivo* study demonstrating lEV mitochondria delivery into motor neurons leave us well-poised to test the hypothesis to rescue motor neuron function in a mouse model of ALS. Overall, neuronal EVs with mitochondrial components represent a promising therapeutic strategy for treating ALS and other neurological disorders.

## Declarations

### Ethics approval and consent to participate

The study does not involve any animal or human data and so, neither ethics approval nor consent to participate was required.

### Consent for publication

*Not applicable (see above)*

### Author Contributions

Conceptualization, Formal analysis, Methodology, Funding acquisition, Project administration, Resources, Supervision D.S.M., O.D., C.M. Investigation, Data curation, Formal analysis, Methodology P.P.P., Z-M.W., P.K., J.R.J., K.S.R., A.L., A.P., K.M.D., S-y. Z., M.C., C.M., O.D., D.S.M. Transmission electron microscopy P.P.P, M.S., D.B.S. *In vitro* assays P.P.P, P.K., J.R.J., K.S.R., A.L., A.P., K.M.D., M.C., S.S.S *In vivo* experiments Z-M.W., O.D., Data analysis P.P.P., C.M., O.D., D.S.M. Manuscript writing P.P.P, P.K., C.M., O.D., D.S.M.

## Data availability

The raw/processed data required to reproduce these findings can be obtained from the corresponding author upon request.

## References

1. Théry, C., et al., Minimal information for studies of extracellular vesicles 2018 (MISEV2018): a position statement of the International Society for Extracellular Vesicles and update of the MISEV2014 guidelines. Journal of Extracellular Vesicles, 2018. 7(1): p. 1535750.

2. Rossi, F.H., M.C. Franco, and A.G. Estevez, Pathophysiology of amyotrophic lateral sclerosis. Current Advances in Amyotrophic Lateral Sclerosis, 2013.

3. Bonafede, R. and R. Mariotti, ALS pathogenesis and therapeutic approaches: the role of mesenchymal stem cells and extracellular vesicles. Frontiers in cellular neuroscience, 2017. 11: p. 80.

4. Goetz, C.G., Amyotrophic lateral sclerosis: early contributions of Jean Martin Charcot. Muscle & Nerve: Official Journal of the American Association of Electrodiagnostic Medicine, 2000. 23(3): p. 336–343.

5. Zarei, S., et al., A comprehensive review of amyotrophic lateral sclerosis. Surgical neurology international, 2015. 6.

6. Bruijn, L.I., T.M. Miller, and D.W. Cleveland, Unraveling the mechanisms involved in motor neuron degeneration in ALS. Annu. Rev. Neurosci., 2004. 27: p. 723–749.

7. Afifi, A., et al., Ultrastructure of atrophic muscle in amyotrophic lateral sclerosis. Neurology, 1966. 16(5): p. 475–475.

8. Atsumi, T., The ultrastructure of intramuscular nerves in amyotrophic lateral sclerosis. Acta neuropathologica, 1981. 55(3): p. 193–198.

9. Sasaki, S., Y. Horie, and M. Iwata, Mitochondrial alterations in dorsal root ganglion cells in sporadic amyotrophic lateral sclerosis. Acta neuropathologica, 2007. 114: p. 633–639.

10. Siklós, L., et al., Ultrastructural evidence for altered calcium in motor nerve terminals in amyotrophc lateral sclerosis. Annals of neurology, 1996. 39(2): p. 203–216.

11. Manfredi, G. and Z. Xu, Mitochondrial dysfunction and its role in motor neuron degeneration in ALS. Mitochondrion, 2005. 5(2): p. 77–87.

12. Jaiswal, M.K., Riluzole but not melatonin ameliorates acute motor neuron degeneration and moderately inhibits SOD1-mediated excitotoxicity induced disrupted mitochondrial Ca2+ signaling in amyotrophic lateral sclerosis. Frontiers in Cellular Neuroscience, 2017. 10: p. 295.

13. Saini, A. and P.A. Chawla, Breaking barriers with tofersen: Enhancing therapeutic opportunities in amyotrophic lateral sclerosis. European Journal of Neurology, 2024. 31(2): p. e16140.

14. Neupane, P., et al., Investigating edaravone use for management of amyotrophic lateral sclerosis (ALS): a narrative review. Cureus, 2023. 15(1).

15. Cha, S.J. and K. Kim, Effects of the edaravone, a drug approved for the treatment of amyotrophic lateral sclerosis, on mitochondrial function and neuroprotection. Antioxidants, 2022. 11(2): p. 195.

16. Fralick, M., C.A. Sacks, and A.S. Kesselheim, Assessment of use of combined dextromethorphan and quinidine in patients with dementia or Parkinson disease after US Food and Drug Administration approval for pseudobulbar affect. JAMA Internal Medicine, 2019. 179(2): p. 224–230.

17. Sun, Y., et al., ALSUntangled# 71: Nuedexta. Amyotrophic Lateral Sclerosis and Frontotemporal Degeneration, 2024. 25(1-2): p. 218–222.

18. 18. FDA-Approved Drugs for Treating ALS. 2024.

19. Zhao, J., et al., The impact of mitochondrial dysfunction in amyotrophic lateral sclerosis. Cells, 2022. 11(13): p. 2049.

20. Cunha-Oliveira, T., et al., Oxidative stress in amyotrophic lateral sclerosis: pathophysiology and opportunities for pharmacological intervention. Oxidative medicine and cellular longevity, 2020. 2020(1): p. 5021694.

21. D’Souza, A., et al., Microvesicles transfer mitochondria and increase mitochondrial function in brain endothelial cells. Journal of Controlled Release, 2021. 338: p. 505–526.

22. Dave, K.M., et al., Engineering extracellular vesicles to modulate their innate mitochondrial load. Cellular and Molecular Bioengineering, 2022. 15(5): p. 367–389.

23. Dave, K.M., et al., Mitochondria-containing extracellular vesicles (EV) reduce mouse brain infarct sizes and EV/HSP27 protect ischemic brain endothelial cultures. Journal of Controlled Release, 2023. 354: p. 368–393

24. Dave, K.M., et al., Mitochondria-containing extracellular vesicles from mouse vs. human brain endothelial cells for ischemic stroke therapy. Journal of Controlled Release, 2024. 373: p. 803–822.

25. Hayakawa, K., et al., Transfer of mitochondria from astrocytes to neurons after stroke. Nature, 2016. 535(7613): p. 551–555.

26. Hayakawa, K., et al., Protective effects of endothelial progenitor cell-derived extracellular mitochondria in brain endothelium. Stem cells, 2018. 36(9): p. 1404–1410.

27. Cselenyák, A., et al., Mesenchymal stem cells rescue cardiomyoblasts from cell death in an in vitro ischemia model via direct cell-to-cell connections. BMC cell biology, 2010. 11: p. 1–11.

28. Masuzawa, A., et al., Transplantation of autologously derived mitochondria protects the heart from ischemia-reperfusion injury. American Journal of Physiology-Heart and Circulatory Physiology, 2013. 304(7): p. H966–H982.

29. Gao, J., et al., Endoplasmic reticulum mediates mitochondrial transfer within the osteocyte dendritic network. Science advances, 2019. 5(11): p. eaaw7215.

30. Liu, D., et al., Intercellular mitochondrial transfer as a means of tissue revitalization. Signal transduction and targeted therapy, 2021. 6(1): p. 65.

31. Torralba, D., F. Baixauli, and F. Sánchez-Madrid, Mitochondria know no boundaries: mechanisms and functions of intercellular mitochondrial transfer. Frontiers in cell and developmental biology, 2016. 4: p. 107.

32. Welsh, J.A., et al., Minimal information for studies of extracellular vesicles (MISEV2023): From basic to advanced approaches. Journal of extracellular vesicles, 2024. 13(2): p. e12404.

33. Todkar, K., et al., Selective packaging of mitochondrial proteins into extracellular vesicles prevents the release of mitochondrial DAMPs. Nature Communications, 2021. 12(1): p. 1971.

34. Guescini, M., et al., Astrocytes and Glioblastoma cells release exosomes carrying mtDNA. Journal of neural transmission, 2010. 117: p. 1–4.

35. Ikeda, G., et al., Mitochondria-rich extracellular vesicles from autologous stem cell– derived cardiomyocytes restore energetics of ischemic myocardium. Journal of the American College of Cardiology, 2021. 77(8): p. 1073–1088.

36. Phinney, D.G., et al., Mesenchymal stem cells use extracellular vesicles to outsource mitophagy and shuttle microRNAs. Nature communications, 2015. 6(1): p. 8472.

37. Puhm, F., et al., Mitochondria are a subset of extracellular vesicles released by activated monocytes and induce type I IFN and TNF responses in endothelial cells. Circulation research, 2019. 125(1): p. 43–52.

38. Liang, W., et al., Mitochondria are secreted in extracellular vesicles when lysosomal function is impaired. Nature Communications, 2023. 14(1): p. 5031.

39. Amari, L. and M. Germain, Mitochondrial extracellular vesicles–origins and roles. Frontiers in Molecular Neuroscience, 2021. 14: p. 767219.

40. Zhou, X., et al., MitoEVs: A new player in multiple disease pathology and treatment. Journal of Extracellular Vesicles, 2023. 12(4): p. 12320.

41. D’Souza, A., et al., Microvesicles transfer mitochondria and increase mitochondrial function in brain endothelial cells. Journal of Controlled Release, 2021. 338: p. 505–526.

42. Dave, K.M., et al., Mitochondria-containing extracellular vesicles (EV) reduce mouse brain infarct sizes and EV/HSP27 protect ischemic brain endothelial cultures. Journal of Controlled Release, 2023. 354: p. 368–393.

43. Islam, M.N., et al., Mitochondrial transfer from bone-marrow–derived stromal cells to pulmonary alveoli protects against acute lung injury. Nature medicine, 2012. 18(5): p. 759–765.

44. O’Brien, C.G., et al., Mitochondria-rich extracellular vesicles rescue patient-specific cardiomyocytes from doxorubicin injury: insights into the SENECA trial. Cardio Oncology, 2021. 3(3): p. 428–440.

45. Jhaveri, J.R., et al., Low pinocytic brain endothelial cells primarily utilize membrane fusion to internalize extracellular vesicles. European Journal of Pharmaceutics and Biopharmaceutics, 2024. 204: p. 114500.

46. Maguire, G., Amyotrophic lateral sclerosis as a protein level, non-genomic disease: Therapy with S2RM exosome released molecules. World Journal of Stem Cells, 2017. 9(11): p. 187.

47. Bonafede, R., et al., Exosome derived from murine adipose-derived stromal cells: neuroprotective effect on in vitro model of amyotrophic lateral sclerosis. Experimental cell research, 2016. 340(1): p. 150–158.

48. Xia, Y., et al., Immunogenicity of extracellular vesicles. Advanced Materials, 2024. 36(33): p. 2403199.

49. Danilushkina, A.A., et al., Strategies for engineering of extracellular vesicles. International journal of molecular sciences, 2023. 24(17): p. 13247.

50. Mouse Motor Neuron-Like Hybrid Cell Line (NSC-34).

51. Madison, R.D., et al., Extracellular vesicles from a muscle cell line (C2C12) enhance cell survival and neurite outgrowth of a motor neuron cell line (NSC-34). Journal of extracellular vesicles, 2014. 3(1): p. 22865.

52. Dave, K.M., et al., Extracellular Vesicles Derived from a Human Brain Endothelial Cell Line Increase Cellular ATP Levels. AAPS PharmSciTech, 2021. 22(1): p. 18.

53. Jhaveri, J.R., et al., Low pinocytic brain endothelial cells primarily utilize membrane fusion to internalize extracellular vesicles. Eur J Pharm Biopharm, 2024. 204: p. 114500.

54. Cottet Rousselle, C., et al., Cytometric assessment of mitochondria using fluorescent probes. Cytometry Part A, 2011. 79(6): p. 405–425.

55. Scharping, N.E., et al., The tumor microenvironment represses T cell mitochondrial biogenesis to drive intratumoral T cell metabolic insufficiency and dysfunction. Immunity, 2016. 45(2): p. 374–388.

56. Maier, O., et al., Differentiated NSC-34 motoneuron-like cells as experimental model for cholinergic neurodegeneration. Neurochemistry international, 2013. 62(8): p. 1029–1038.

57. Cashman, N.R., et al., Neuroblastoma× spinal cord (NSC) hybrid cell lines resemble developing motor neurons. Developmental dynamics, 1992. 194(3): p. 209–221.

58. Kaur, R., et al., Recent developments in tubulin polymerization inhibitors: an overview. European journal of medicinal chemistry, 2014. 87: p. 89–124.

59. Jouhilahti, E.-M., S. Peltonen, and J. Peltonen, Class III β-tubulin is a component of the mitotic spindle in multiple cell types. Journal of Histochemistry & Cytochemistry, 2008. 56(12): p. 1113–1119.

60. Doyle, L.M. and M.Z. Wang, Overview of extracellular vesicles, their origin, composition, purpose, and methods for exosome isolation and analysis. Cells, 2019. 8(7): p. 727.

61. Morad, G., et al., Tumor-derived extracellular vesicles breach the intact blood–brain barrier via transcytosis. ACS nano, 2019. 13(12): p. 13853–13865.

62. Raposo, G. and W. Stoorvogel, Extracellular vesicles: exosomes, microvesicles, and friends. Journal of Cell Biology, 2013. 200(4): p. 373–383.

63. Woo, H.-K., et al., Characterization and modulation of surface charges to enhance extracellular vesicle isolation in plasma. Theranostics, 2022. 12(5): p. 1988.

64. van Niel, G., G. D’Angelo, and G. Raposo, Shedding light on the cell biology of extracellular vesicles. Nature Reviews Molecular Cell Biology, 2018. 19(4): p. 213–228.

65. Arif, S. and V.J. Moulin, Extracellular vesicles on the move: Traversing the complex matrix of tissues. European journal of cell biology, 2023. 102(4): p. 151372.

66. Han, M., et al., Transfection study using multicellular tumor spheroids for screening non-viral polymeric gene vectors with low cytotoxicity and high transfection efficiencies. Journal of Controlled Release, 2007. 121(1-2): p. 38–48.

67. Jung, M.K. and J.Y. Mun, Sample preparation and imaging of exosomes by transmission electron microscopy. Journal of visualized experiments: JoVE, 2018(131).

68. Gheldof, D., et al., Thrombin generation assay and transmission electron microscopy: a useful combination to study tissue factor-bearing microvesicles. Journal of extracellular vesicles, 2013. 2(1): p. 19728.

69. Vechetti Jr, I.J., Emerging role of extracellular vesicles in the regulation of skeletal muscle adaptation. Journal of Applied Physiology, 2019. 127(2): p. 645–653.

70. Jeppesen, D.K., et al., Extracellular vesicles and nanoparticles: emerging complexities. Trends in Cell Biology, 2023. 33(8): p. 667–681.

71. Bobryshev, Y.V., M.C. Killingsworth, and A.N. Orekhov, Increased shedding of microvesicles from intimal smooth muscle cells in athero-prone areas of the human aorta: implications for understanding of the predisease stage. Pathobiology, 2012. 80(1): p. 24–31.

72. Dave, K.M., D.B. Stolz, and D.S. Manickam, Delivery of mitochondria-containing extracellular vesicles to the BBB for ischemic stroke therapy. Expert Opinion on Drug Delivery, 2023. 20(12): p. 1769–1788.

73. Haney, M.J., et al., Extracellular vesicles as drug delivery system for the treatment of neurodegenerative disorders: Optimization of the cell source. Advanced nanobiomed research, 2021. 1(12): p. 2100064.

74. Tognoli, M.L., et al., Lack of involvement of CD63 and CD9 tetraspanins in the extracellular vesicle content delivery process. Communications Biology, 2023. 6(1): p. 532.

75. Dozio, V. and J.-C. Sanchez, Characterisation of extracellular vesicle-subsets derived from brain endothelial cells and analysis of their protein cargo modulation after TNF exposure. Journal of Extracellular Vesicles, 2017. 6(1): p. 1302705.

76. Wang, W., et al., Biogenesis and function of extracellular vesicles in pathophysiological processes of skeletal muscle atrophy. Biochemical pharmacology, 2022. 198: p. 114954.

77. Yokoi, A. and T. Ochiya. Exosomes and extracellular vesicles: Rethinking the essential values in cancer biology. in Seminars in cancer biology. 2021. Elsevier.

78. Kumar, M.A., et al., Extracellular vesicles as tools and targets in therapy for diseases. Signal transduction and targeted therapy, 2024. 9(1): p. 27.

79. Ikeda, G., et al., Mitochondria containing extracellular vesicles from autologous induced pluripotent stem cell derived cardiomyocytes restore bioenergetics in ischemic myocardium. Journal of the American College of Cardiology, 2020. 75(11_Supplement_1): p. 3659–3659.

80. Hayakawa, K., et al., Transfer of mitochondria from astrocytes to neurons after stroke. Nature, 2016. 535: p. 551.

81. Ikeda, G., et al., Mitochondria-Rich Extracellular Vesicles From Autologous Stem Cell-Derived Cardiomyocytes Restore Energetics of Ischemic Myocardium. J Am Coll Cardiol, 2021. 77(8): p. 1073–1088.

82. Phinney, D.G., et al., Mesenchymal stem cells use extracellular vesicles to outsource mitophagy and shuttle microRNAs. Nature Communications, 2015. 6: p. 8472.

83. Rikkert, L.G., et al., Quality of extracellular vesicle images by transmission electron microscopy is operator and protocol dependent. Journal of extracellular vesicles, 2019. 8(1): p. 1555419.

84. Dave, K.M., et al., Mitochondria-containing extracellular vesicles from mouse vs. human brain endothelial cells for ischemic stroke therapy. bioRxiv, 2024: p. 2024.01. 16.575903.

85. Robinson, M.B., et al., Exogenous Hsc70, but not thermal preconditioning, confers protection to motoneurons subjected to oxidative stress. Developmental Neurobiology, 2008. 68(1): p. 1–17.

86. Habte, M.L. and E.A. Beyene, Biological application and disease of oxidoreductase enzymes, in Oxidoreductase. 2020, IntechOpen.

87. Guescini, M., et al., C2C12 myoblasts release micro-vesicles containing mtDNA and proteins involved in signal transduction. Experimental cell research, 2010. 316(12): p. 1977–1984.

88. Sansone, P., et al., Packaging and transfer of mitochondrial DNA via exosomes regulate escape from dormancy in hormonal therapy-resistant breast cancer. Proceedings of the National Academy of Sciences, 2017. 114(43): p. E9066–E9075.

89. Théry, C., et al., Molecular characterization of dendritic cell-derived exosomes: selective accumulation of the heat shock protein hsc73. The Journal of cell biology, 1999. 147(3): p. 599–610.

90. Shi, C., et al., Characterization of heat shock protein 27 in extracellular vesicles: a potential anti inflammatory therapy. The FASEB Journal, 2019. 33(2): p. 1617–1630.

91. Braganza, A., G.K. Annarapu, and S. Shiva, Blood-based bioenergetics: An emerging translational and clinical tool. Mol Aspects Med, 2020. 71: p. 100835.

92. Nolfi-Donegan, D., A. Braganza, and S. Shiva, Mitochondrial electron transport chain: Oxidative phosphorylation, oxidant production, and methods of measurement. Redox Biol, 2020. 37: p. 101674.

93. Sure, V.N., et al., A novel high-throughput assay for respiration in isolated brain microvessels reveals impaired mitochondrial function in the aged mice. Geroscience, 2018. 40(4): p. 365–375.

94. Braganza, A., et al., Platelet bioenergetics correlate with muscle energetics and are altered in older adults. JCI insight, 2019. 5(13): p. e128248.

95. Cardenes, N., et al., Platelet bioenergetic screen in sickle cell patients reveals mitochondrial complex V inhibition, which contributes to platelet activation. Blood, 2014. 123(18): p. 2864–2872.

96. Gu, X., et al., Measurement of mitochondrial respiration in adherent cells by Seahorse XF96 Cell Mito Stress Test. STAR protocols, 2021. 2(1): p. 100245.

97. Dave, K.M., et al., Mitochondria-containing extracellular vesicles from mouse vs. human brain endothelial cells for ischemic stroke therapy. Journal of Controlled Release, 2024. 373: p. 803–822.

98. Puhm, F., Mitochondria Are a Subset of Extracellular Vesicles Released by Activated Monocytes and Induce Type I IFN and TNF Responses in Endothelial Cells (vol 125, pg 43, 2019). CIRCULATION RESEARCH, 2019. 125(10): p. E93–E93.

99. Xia, L., et al., AdMSC-derived exosomes alleviate acute lung injury via transferring mitochondrial component to improve homeostasis of alveolar macrophages. Theranostics, 2022. 12(6): p. 2928.

