## Supplemental File for "Mitochondria-containing large extracellular vesicles target mouse motor neurons upon intramuscular injection"

**Supplementary information**

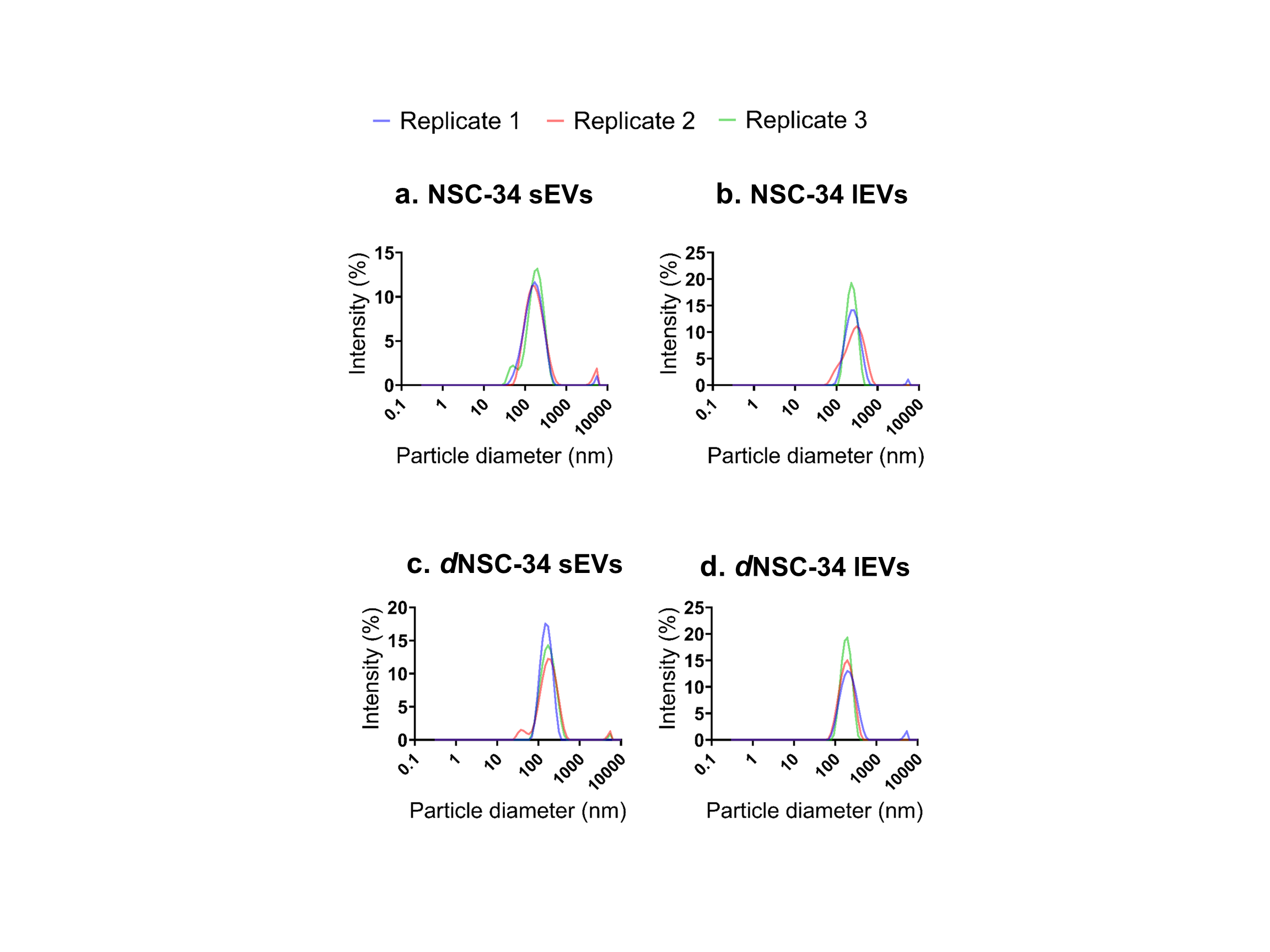

**Supplementary Figure 1.** Intensity-weight particle size distribution of **a.** NSC-34 sEVs**; b.** NSC-34 lEVs**; c.** *d*NSC-34 sEVs; **d.** *d*NSC-34 lEVs using dynamic light scattering demonstrate the distribution of particle diameter as a function of % scattered light intensity.

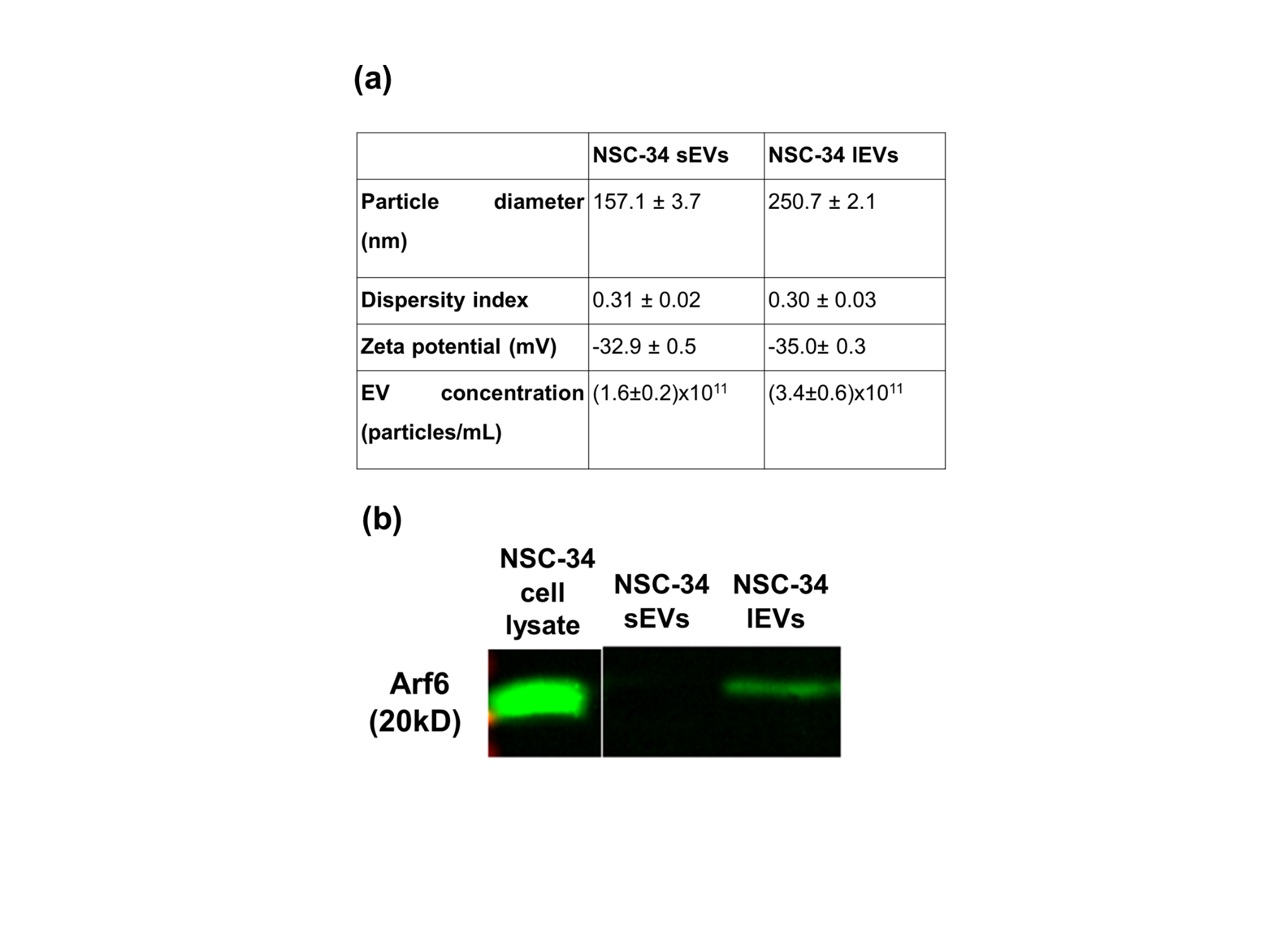

**Supplementary Figure 2. Physicochemical characterization and western blotting of NSC-34 sEVs and lEVs. (a)** NSC-34-derived sEVs and lEVs were characterized by measuring their particle diameter, dispersity indices, and zeta potential using DLS, Malvern Zetasizer Pro (Malvern Panalytical Inc., Westborough, PA). Samples were diluted at a 0.1 mg/ml concentration using either PBS or 10 mM HEPES at pH 7.4 to measure particle size and zeta potential, respectively. Samples were run in triplicate and data were represented as mean± standard deviation (SD). **(b)** Western blotting of NSC-34 derived sEVs and lEVs to detect protein (Arf6) involved in EV biogenesis. Western blotting showed expression of Arf6 in lEVs not in sEVs. The samples were electrophoresed using sodium dodecyl sulfate-polyacrylamide gel followed by transfer to a PVDF membrane. After treatment with blocking buffer, samples were incubated overnight with primary antibody at 4^0^C and with secondary antibody at room temperature. The blot was imaged using Odyssey imager (LI-COR Inc. Lincoln, NE) under an 800 nm 700nm channel.

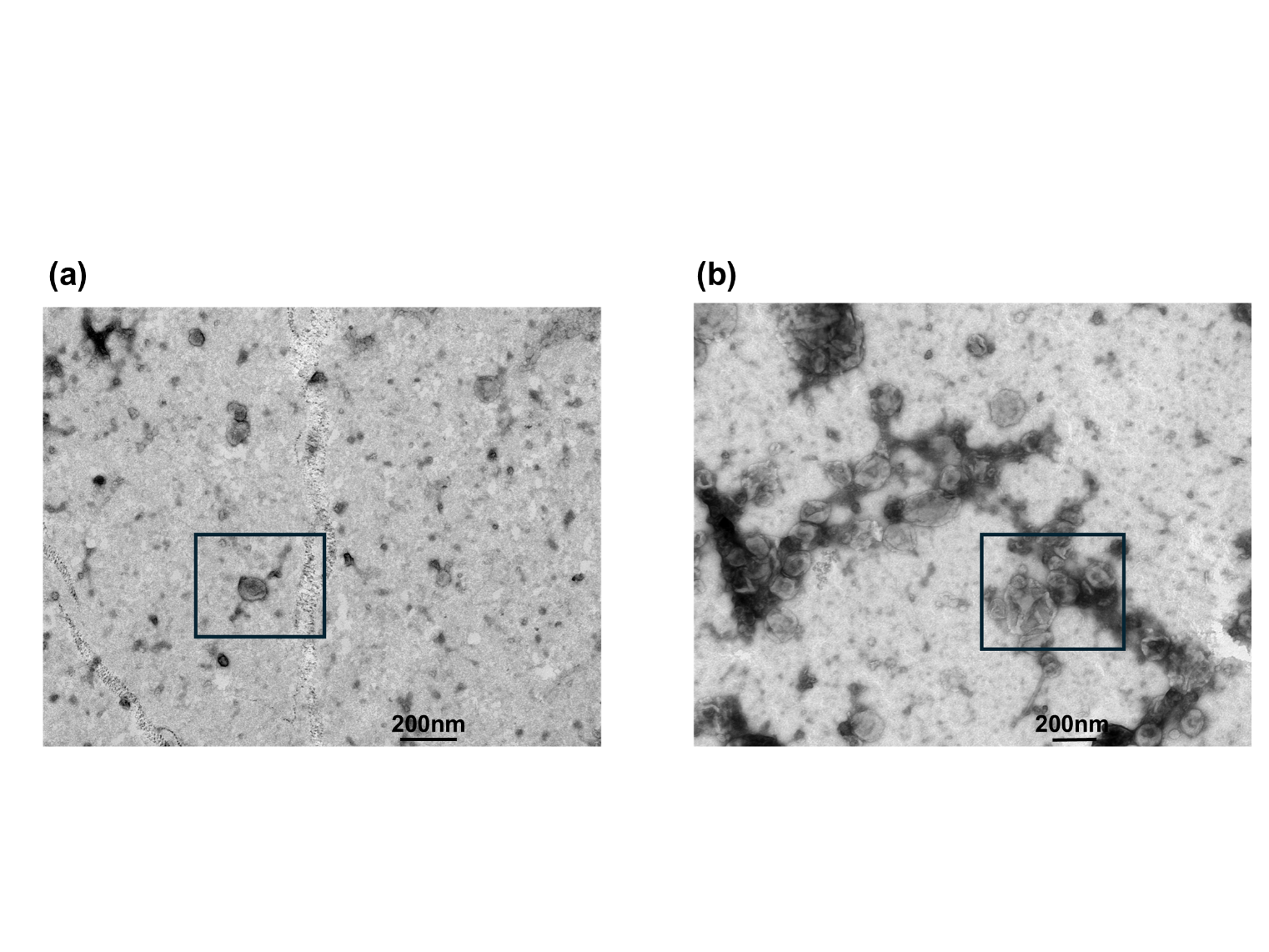

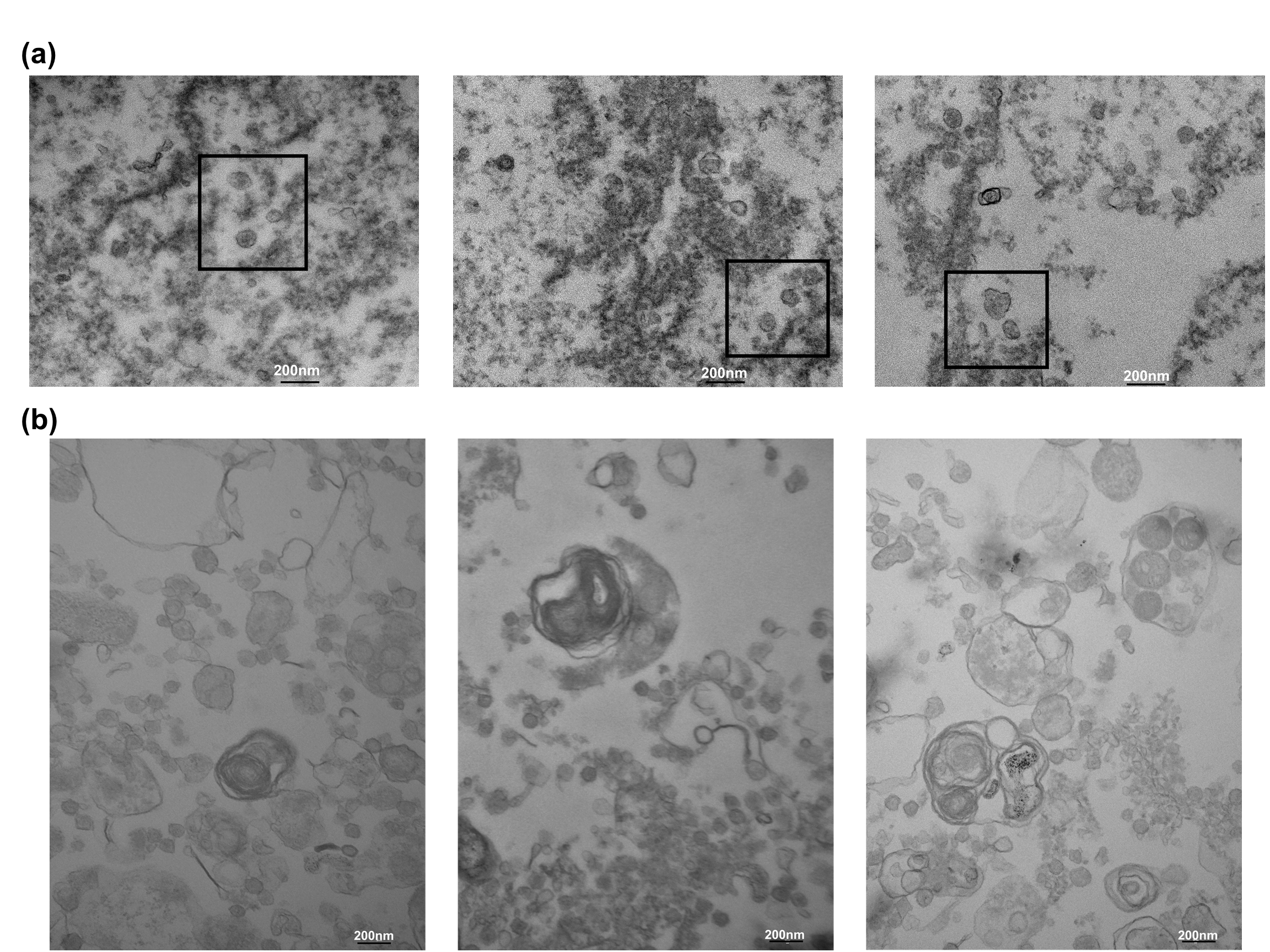
**Supplementary Figure 3.** Negative stain TEM image of *d*NSC-34-derived **a.** sEVs and **b.** lEVs. Scale bar: 200 nm

**Supplementary Figure 4.** Cross-sectioned TEM images of *d*NSC-34-derived **a.** sEVs and **b.** m/lEVs. Scale bar: 200 nm.

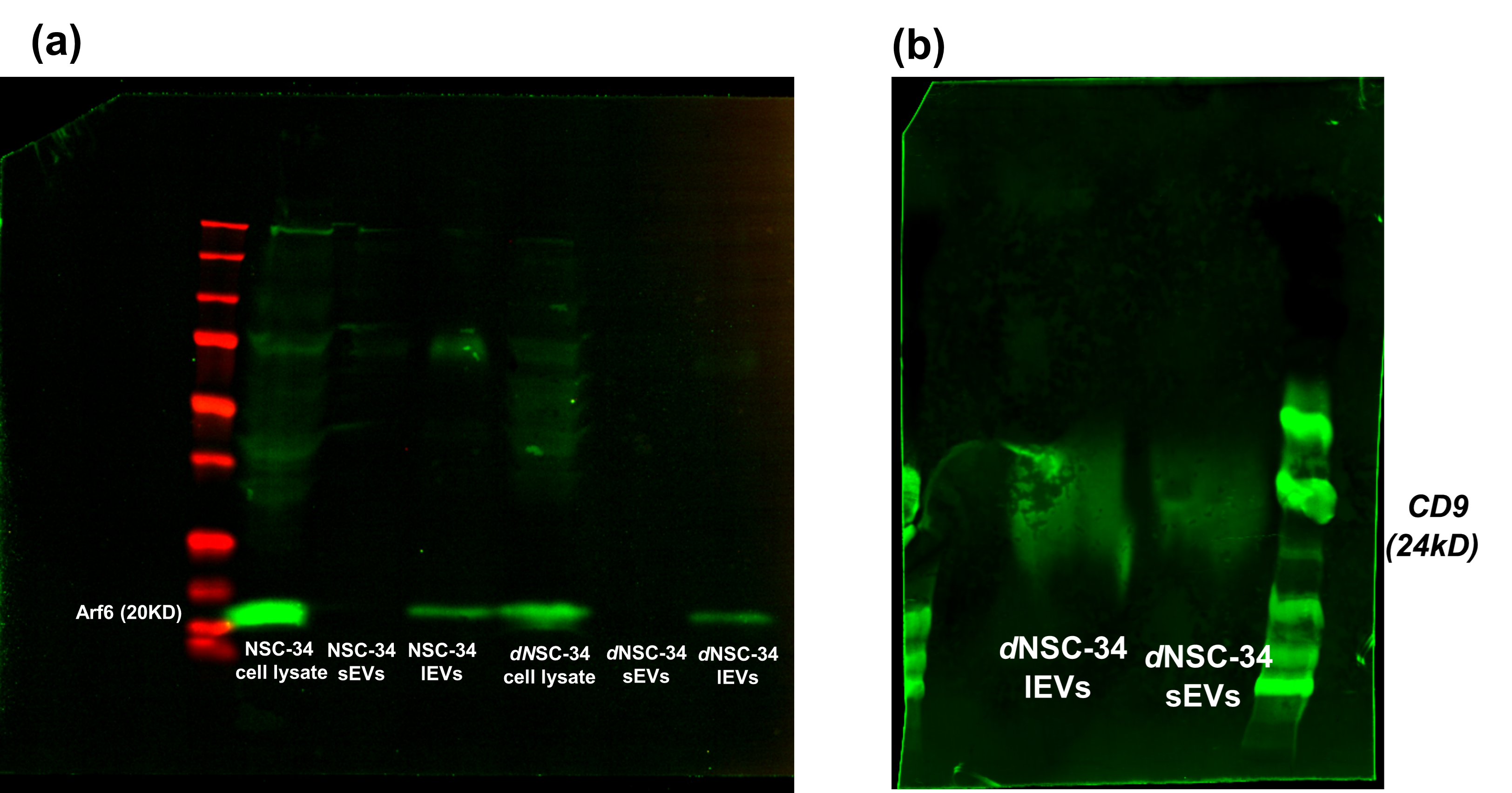

**Supplementary Figure 5.** Western blotting of *d*NSC-34-derived EVs and NSC-34-derived EVs. Western blotting to detect Arf6 protein involved in lEV biogenesis and EV marker protein CD9. The samples were electrophoresed using sodium dodecyl sulfate-polyacrylamide gel followed by transfer to a PVDF membrane. After treatment with blocking buffer, samples were incubated overnight with primary antibody at 4^0^C and with secondary antibody at room temperature. The blot was imaged using the Odyssey imager (LI-COR Inc. Lincoln, NE) under an 800 nm 700nm channel.

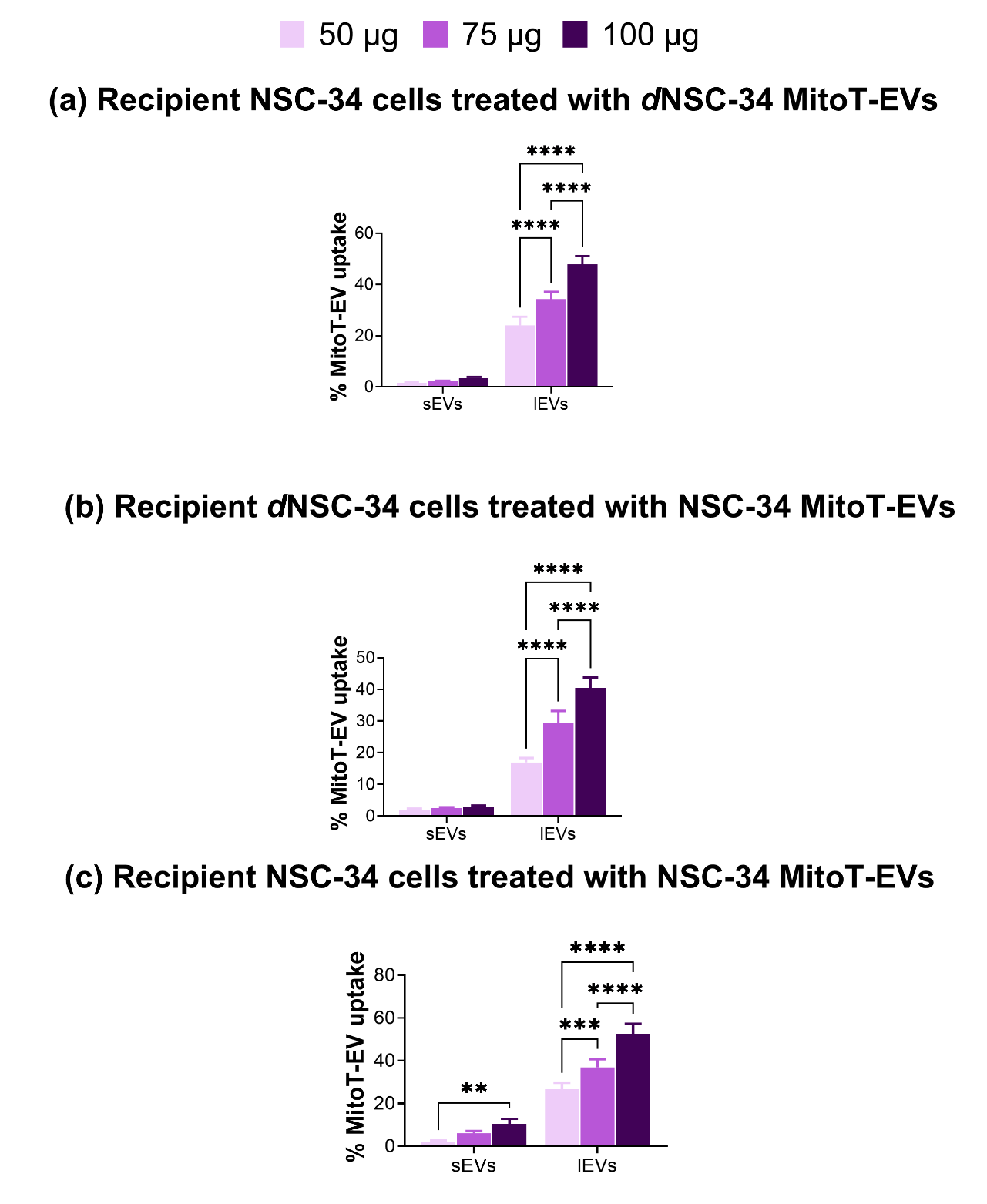

**Supplementary Figure 6. Quantification of EV-mediated mitochondrial transfer into recipient cells.** The signal intensity of MitoT-red in recipient cells obtained from flow cytometry was quantified indicating EV-mediated mitochondrial transfer (**a)** Recipient NSC-34 cells treated with indicated doses of *d*NSC-34-derived MitoT-red labeled lEVs or sEVs**. (b)** Recipient *d*NSC-34 cells treated with indicated doses of NSC-34 derived MitoT-red labeled lEVs or sEVs. (**c)** Recipient NSC-34 cells were treated with indicated doses of NSC-34 MitoT-lEVs or MitoT-sEVs. Data represented as mean ± SD of n=4.

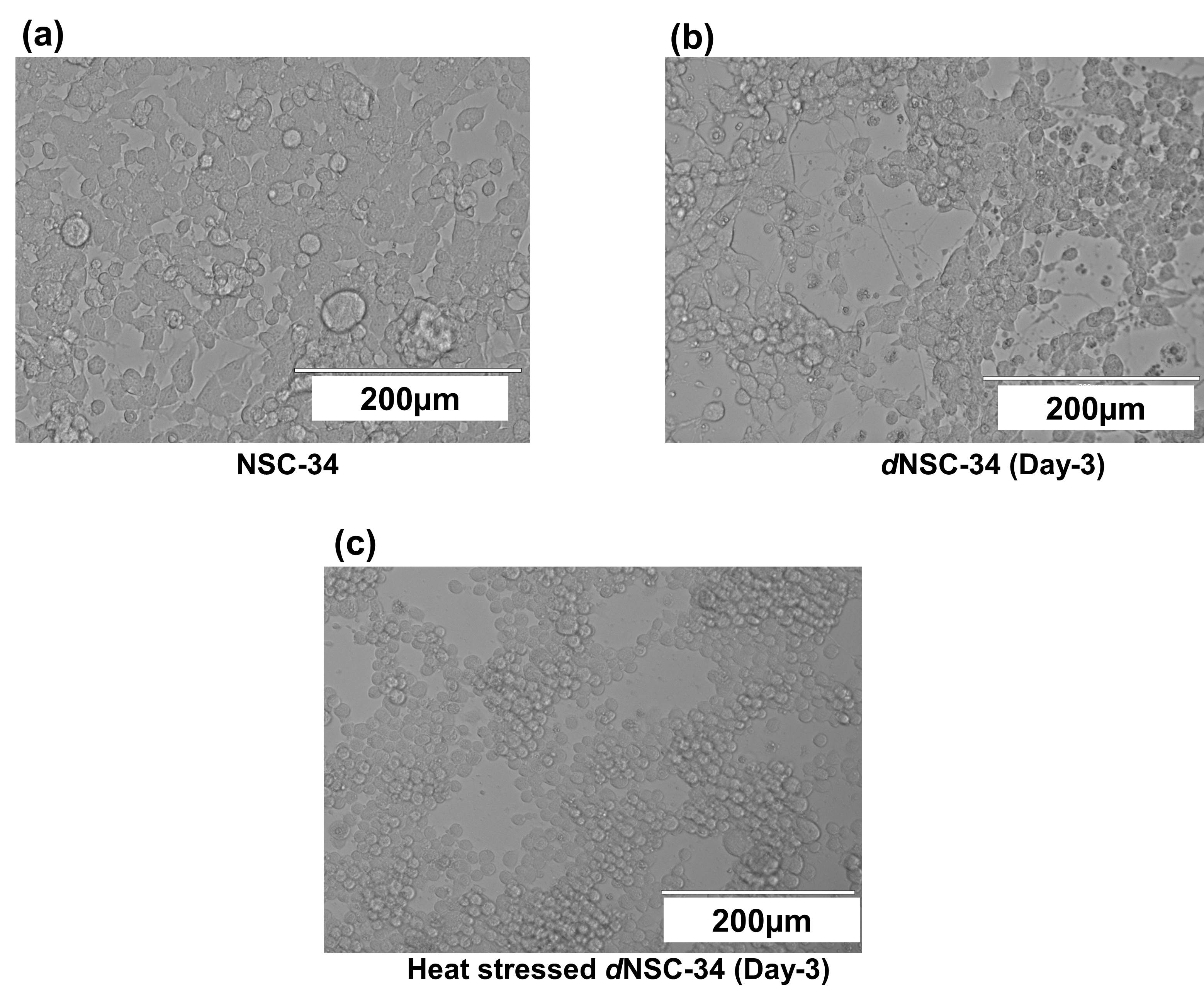

**Supplementary Figure 7. EVOS microscopic images of heat-stressed *d*NSC-34 cells (neurons).** On day 3 of differentiation, *d*NSC-34 cells were treated with Neurobasal differentiation medium pre-heated at 50.8⁰C for 30 minutes followed by additional 30 minutes incubation at 50.8⁰C. Heat stressed *d*NSC-34 cells or neurons were observed under EVOS microscope to study the morphological changes as an impact of heat stress. (**a)** undifferentiated NSC-34 grown with DMEM high glucose medium, 10 % FBS, and 1% pen-strep. (**b)** Untreated healthy *d*NSC-34 cells. **(c)** *d*NSC-34 (neurons) under heat stress showed loss of neurite outgrowth. All images were captured at 20X magnification under the transmitted channel. Scale bar: 200 µm.
